# Megabase-scale loss of heterozygosity provoked by CRISPR-Cas9 DNA double-strand breaks

**DOI:** 10.1101/2024.09.27.615517

**Authors:** Samantha B. Regan, Darpan Medhi, Travis B. White, Yi-Zhen Jiang, Su Jia, Qichen Deng, Maria Jasin

## Abstract

Harnessing DNA double-strand breaks (DSBs) is a powerful approach for gene editing, but it may provoke loss of heterozygosity (LOH), which predisposes to tumorigenesis. To interrogate this risk, we developed a two-color flow cytometry-based system (Flo-LOH), detecting LOH in ∼5% of cells following a DSB. After this initial increase, cells with LOH decrease due to a competitive disadvantage with parental cells, but if isolated, they stably propagate. Segmental loss from terminal deletions with de novo telomere addition and nonreciprocal translocations is observed as well as whole chromosome loss, especially following a centromeric DSB. LOH spans megabases distal from the DSB, but also frequently tens of megabases centromere-proximal. Inhibition of microhomology-mediated end joining massively increases LOH, which is synergistically increased with concomitant inhibition of canonical nonhomologous end joining. The capacity for large-scale LOH must therefore be considered when using DSB-based gene editing, especially in conjunction with end joining inhibition.

## Introduction

Loss of heterozygosity (LOH), which refers to the loss of genetic information from one parental genome, is a frequent occurrence in cancer cells. LOH can arise through several mechanisms, including large-scale changes in copy number, such as from chromosome truncation. Such LOH events are deleterious because they cause protein imbalances and advance genomic instability^1^. Copy neutral LOH, such as that which arises from crossing over during interhomolog homologous recombination (IH-HR) or whole chromosome loss with duplication of the remaining chromosome, is also problematic. With copy neutral LOH in cells heterozygous for a mutation in a tumor suppressor gene, the wild-type copy of the gene can be replaced by the mutant copy, which is an initiating event for tumorigenesis^2,3^. Thus, LOH is prevalent in cancer cells because it can be an initial driver of the disease and, in cells with ongoing genomic instability, shape tumor evolution to promote cell proliferation and metastasis^4–7^. On the flip side, LOH can have beneficial effects in patients when it leads to loss of dominant mutations^8^.

DNA double-strand breaks (DSBs) are likely initiating lesions in many instances of LOH. Genome editors such as CRISPR-Cas9 are powerful tools for altering the genome through the introduction of DSBs, with widespread use in cultured cells, model organisms, and, more recently, patients. While Cas9-targeted DSBs typically result in deletions or insertions of a few bps, disrupting gene function^9^, moderately-sized deletions in the kilobase to megabase range are not infrequent after a Cas9 DSB^10–20^. However, investigations in recent years have identified instances of even more significant LOH with profound consequences to the genome. For example, major genomic changes involving chromosome truncations and loss of whole chromosomes have been identified to be common in human embryos after introduction of a Cas9 DSB^14,21^, questioning the use of genome editing involving targeted DSBs in embryos. Recently, large structural changes have been found in other systems as well^22–25^, including primary human T cells^26,27^, which are employed in somatic gene editing for cancer treatment. In addition to considerations in clinical settings, LOH from chromosomal truncations may complicate conclusions drawn from CRISPR screens^28^. Copy neutral LOH, with megabases of large-scale LOH but no deletions, has also been observed after a DSB in several models^18,20,21,24,29–32^. Included among these are chromothriptic events in which copy number is maintained but segments are erratically rearranged^23^.

Given its cellular impact, we sought to interrogate causes of LOH and factors that impact it. However, analysis of LOH can often be laborious and expensive. With the goal of streamlining this process, we established a two-color flow cytometry-based system we termed Flo-LOH in mouse embryonic stem cells, which allows for efficient single-cell quantification and isolation of cells that have undergone long tracts of LOH. We find that LOH typically occurs in several percent of cells after a DSB. Although cells with LOH are outcompeted by heterozygous cells over time, they survive and proliferate when isolated from the population.

Extensive characterization of isolated cells uncovered both segmental LOH and whole chromosome loss, the latter of which is especially common with a DSB close to the centromere. Segmental LOH frequently involves terminal deletions from the DSB site and, to a lesser extent, nonreciprocal translocations. Surprisingly, tracts of LOH often extend megabases centromere-proximal from the DSB, as well as distal. Inhibition of end joining pathways profoundly increases LOH. While LOH from IH-HR is infrequent, Flo-LOH identifies rare IH-HR events that lead to LOH. Finally, we demonstrate the power of Flo-LOH as a sensor of general genomic instability when the DNA repair factor BLM is depleted, or DNA is damaged globally by radiation.

## Results

### An allele-specific DSB leads to frequent loss of heterozygosity

Given the impact of LOH on cells, we developed a two-color system to measure LOH by flow cytometry, termed Flo-LOH (**Fig. 1A**). Fluorescent markers *mRuby3* and *dClover2* were targeted to the two alleles of the *Klf5* gene on chromosome 14 (chr14) in mouse embryonic stem cells (mESCs) using CRISPR- Cas9 (**Fig. S1A,B**). The markers were fused to the 3’ end of *Klf5* with an intervening T2A peptide sequence, such that fluorescent marker expression was driven by the endogenous *Klf5* promoter without creating a fusion protein. To have polymorphisms between chromosome homologs, the mESCs are derived from F1 hybrid mice, with one set of chromosomes from the 129/Sv strain (129) and the other from the C57BL/6 strain (B6)^33^. Four double-positive clones were used for this analysis, two for each chr14 orientation of the fluorescent markers, i.e., *mRuby3*^129^*;dClover2*^B6^ and *dClover2*^129^;*mRuby3* ^B6^. In all such *Klf5^dClover^*^2^*^/mRuby^*^3^ clones, bright dClover2 and mRuby3 fluorescence is observed in >99% of cells (**Fig. 1B, S1C**). The few remaining cells (∼0.4%) express just one fluorescent marker, possibly due to spontaneous LOH.

**Figure 1.**
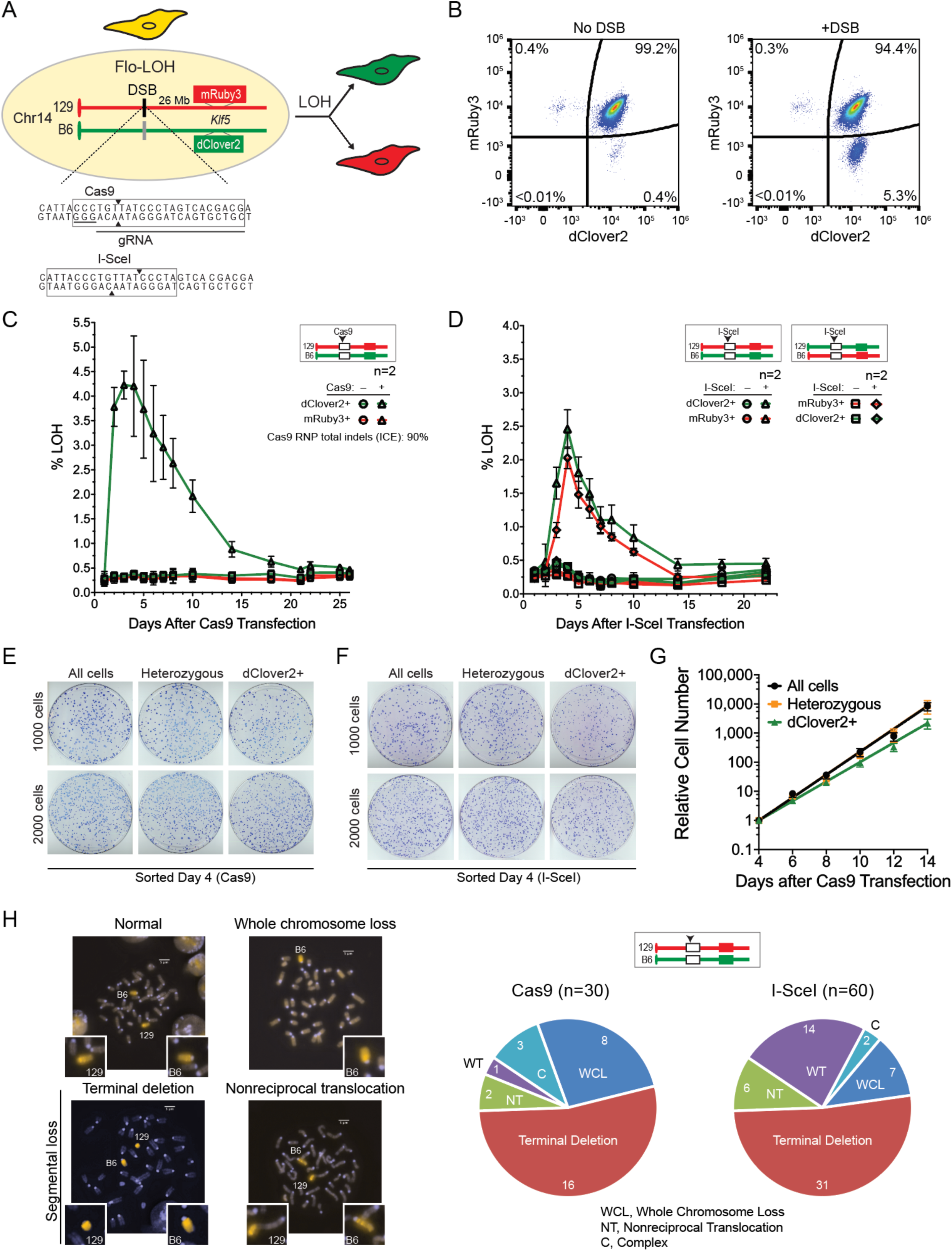
A DSB on a single chromosome leads to frequent loss of heterozygosity. **A.** Flo-LOH. Fluorescent markers *dClover2* and *mRuby3* coding sequences were knocked into each chr14 of 129/B6 F1 hybrid mouse embryonic stem cells (mESCs) at the *Klf5* locus. Fluorescent markers are expressed from the *Klf5* promoter but separated from the protein by a T2A self-cleaving peptide sequence. As shown, a DSB is specifically introduced into the 129 chr14 by either I-SceI or Cas9 at overlapping recognition sites. The Cas9 PAM is underlined. Cas9 makes a blunt-ended DSB, while I-SceI gives rise to a 4 base 3’ overhang. **B.** Spontaneous LOH is low, but a DSB leads to loss of the fluorescent marker expressed from the broken chromosome. Prior to the DSB, >99% of cells express both dClover2 and mRuby3 as seen by flow cytometry. Induction of a DSB specifically on the 129 chr14 leads to ∼5% single-positive dClover2 cells. **C,D.** Time course of LOH after induction of a DSB on the 129 chr14 in Flo-LOH cell lines. The percentage of cells with LOH in the population peaks within 3-4 days after DSB induction by a Cas9 RNP (**C**) or I-SceI plasmid expression (**D**), but then decreases over time. I-SceI experiments were carried out in both chromosomal orientations of the fluorescent markers, with reciprocal results (**D**). n=2 for each condition with 3 plus DSB replicates within each experiment and one minus DSB replicate. **E,F.** Cells with LOH form colonies after DSB induction at a slightly reduced frequency compared to those without LOH. Cells were sorted by flow cytometry four days after transfection with a Cas9 RNP (**B**) or an I-SceI expression vector (**D**) and plated at the indicated number of cells to form colonies. (See also **Fig. S2B**.) **G.** Cells with LOH proliferate ∼15% more slowly than heterozygous cells. Cells were sorted four days after Cas9 RNP transfection and 50,000 were plated. Every 48 hr cells were recounted and 50,000 were replated. One representative experiment is shown. (See also **Fig. S2C**.) **H.** DSB induction leads to chromosome aberrations. After DSB induction by Cas9 or I-SceI, single-positive dClover2 cell populations were sorted, and metaphase spreads were stained with DAPI and probed with a whole chr14 probe. Terminal deletions were observed in the majority of cells with aberrations, but whole chromosome loss and nonreciprocal translocations were also observed at lower frequency. Left, representative images of aberrations; Right, quantification of types of aberrations.

To determine the impact of a single double-strand break (DSB) on chromosome integrity, we introduced either Cas9 or I-SceI endonuclease to specifically cleave the 129 chr14 (**Fig. 1A**). The DSB is located 26 Mb centromere-proximal to the *mRuby3* fluorescent marker, such that loss of mRuby3 expression after DSB formation would entail a large-scale genomic event. After delivering Cas9-gRNA as a ribonucleoprotein particle (RNP), >4% of cells had lost expression of mRuby3 (**Fig. 1B,C**). Cas9 cleavage was highly efficient as estimated by ICE analysis of indels (90%), implying that LOH events, while significant in number, were a fraction of total repair events. The proportion of single-positive dClover2 cells peaked 3 to 4 days after RNP transfection, but then decreased to nearly background levels by ∼3 weeks (**Fig. 1C**).

Expressing I-SceI from a plasmid, which is somewhat less efficient, also led to a large initial increase in single-positive dClover2 cells, which then decreased (**Fig. 1D** **and S2A**). Switching the chromosomal orientation of the fluorescent markers gave reciprocal results, with a large, transient increase in single-positive mRuby3 cells (**Fig. 1D**).

The decrease in single-positive cells over time could be the result of cell lethality and/or a growth disadvantage. To examine these possibilities, single-positive dClover2 cells were sorted 4 days after transfection and plated to form single colonies. Colonies readily formed from dClover2+ cells, although at a slightly reduced frequency compared to sorted heterozygous cells (**Fig. 1E,F and S2B**). Additionally, the doubling time of dClover2+ cells was ∼15% longer than heterozygous cells (**Fig. 1G and S2C**). These results imply that the LOH in this context causes minimal lethality and a slight growth disadvantage that becomes pronounced when cells are grown in competition with >90% heterozygous cells.

### Segmental loss of >50 Mb is common after a DSB

To determine the types of chromosomal changes that led to LOH, we sorted single-positive dClover2 cells 4 days after transfection with either the Cas9 RNP or the I-SceI expression vector and performed fluorescence in situ hybridization (FISH) on the cell populations with whole chr14 paints. Fortuitously, the 129 and B6 chr14s could be distinguished by differential DAPI staining of the centromeres (**Fig. S2D**). Most sorted LOH cells had an observable aberration on the 129 chr14, predominantly a terminal deletion (**Fig. 1H**). Less frequently, nonreciprocal translocation with loss of the chr14 region distal to the DSB was also observed, as was whole chromosome loss, which occurred with or without duplication of the B6 chr14 (12 and 3 metaphase spreads, respectively). Thus, loss of mRuby3 fluorescence resulted from LOH which minimally extends 52 Mb distal from the DSB site and maximally encompasses the entire chromosome.

To further characterize LOH events, we single-cell sorted individual single-positive dClover2 cells on both day 4 and day 10 after transfection (**Fig. 2A**). Cells readily expanded to form colonies and maintained expression of dClover2 only. The *mRuby3* gene, located 26 Mb from the DSB, was absent in all but two clones. The *mRuby3* coding sequence was mutated in one clone, but unaltered in the other; these two clones were eliminated from further analysis. The distal *D14Mit95* polymorphism, located 37 Mb from the DSB, had undergone LOH in all of the clones, such that only the B6 allele was maintained, supporting that these clones had terminal deletions rather than interstitial deletions that extend to the fluorescent marker.

**Figure 2.**
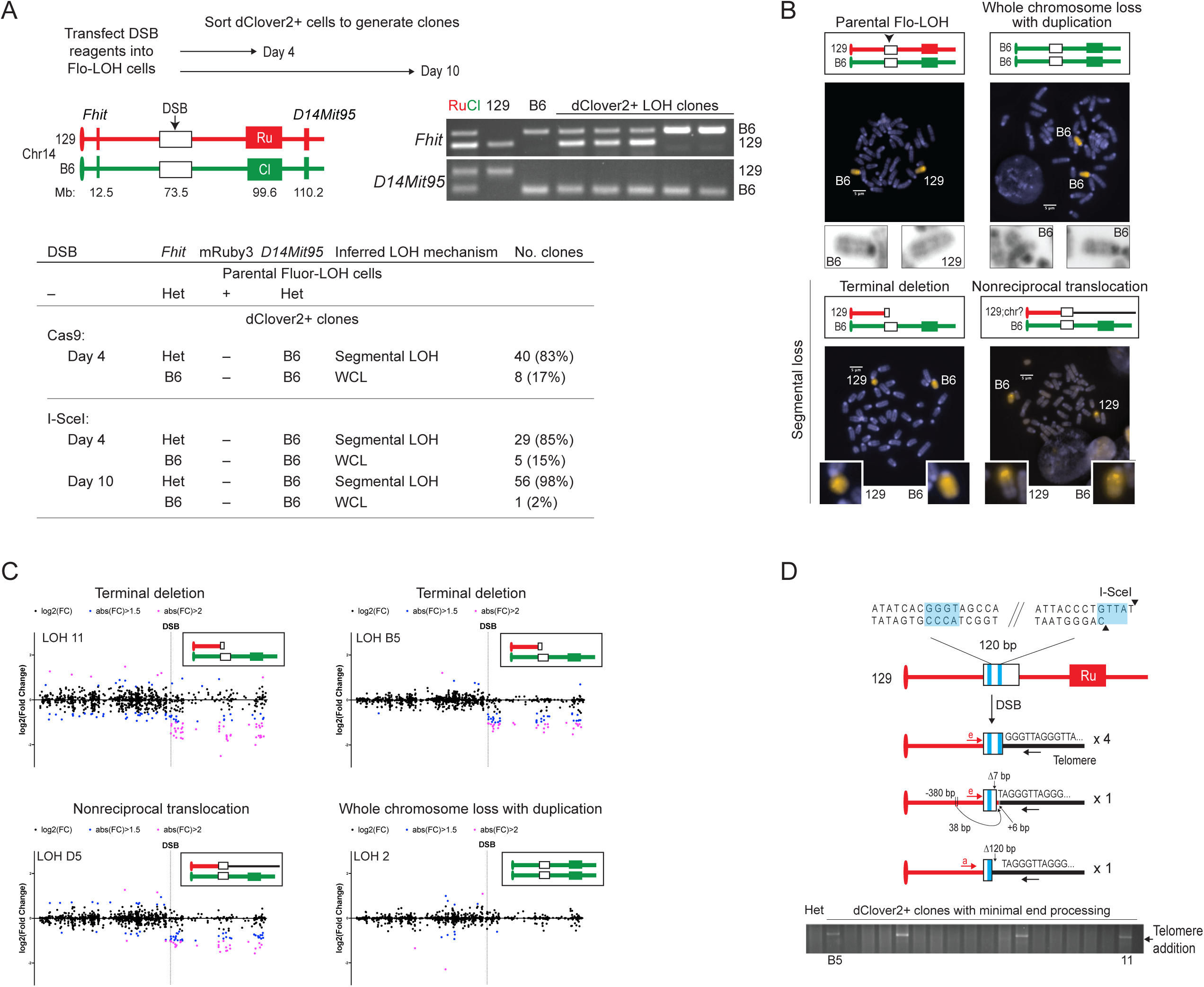
Mid-chromosome breakage leads to frequent segmental loss. **A.** Segmental loss is frequent in cells with LOH after a DSB. Single-positive dClover2 cells were sorted 4 or 10 days after transfection with a Cas9 RNP or an I-SceI expression vector to introduce a DSB on the 129 chr14. Clones were analyzed at a centromere-proximal *Fhit* polymorphism and the distal *D14Mit95* polymorphism. Segmental loss is detected in the vast majority clones through retention of only the B6 *D14Mit95* allele but maintenance of heterozygosity at the *Fhit* polymorphism. Whole chromosome loss is inferred in a smaller fraction of clones from retention of only the B6 alleles at both *Fhit* and *D14Mit95*. RuCl, DNA from Flo-LOH parental cells; 129 and B6, DNA from 129 cells and B6 cells, respectively; WCL, whole chromosome loss. **B.** Whole chromosome painting with a chr14 probe confirms genomic loss in the dClover2+ clones. The B6 chromosome and 129 chromosomes can be differentiated by the intensity of DAPI staining of the centromere. (See also **Fig. S2D**.) Segmental loss occurs from either terminal deletion or nonreciprocal translocation. Whole chromosome (129 chr14) loss in one clone was associated with duplication of the B6 chr14. Clones are derived from the day 10 sort. **C.** Decreased expression of genes distal to the DSB on chr14 in clones with segmental loss, as determined by RNA-seq analysis, but not in the clone with whole chromosome loss with duplication. (See also **Fig. S3**.) **D.** *De novo* telomere addition in clones with terminal deletion. PCR was performed on dClover2+ clones using a forward primer (a or e) near the I-SceI site on chr14 and a reverse primer binding to telomere repeats. In 4 clones, telomere addition occurred at 4 bp in the I-SceI site that have microhomology to a telomere repeat, whereas in another clone it occurred at 4 bp of microhomology to the telomere repeat located 120 bp upstream. (Four base identities in blue shading.) Telomere addition in the remaining clone was more complicated, involving a 7 bp deletion at the I-SceI site, templated insertion of a 38 bp sequence located ∼380 bp distal to the I-SceI site, and another 6 bp insertion. Below, gel of telomere addition PCR in day 10 clones that have minimal end processing (HindIII site PCR, **Fig. S4A-C**). Labelled clones LOH B5 and LOH 11 with telomere addition have terminal deletions and were included in the RNA-Seq analysis (Fig. 2C).

In most clones, the centromere-proximal *Fhit* polymorphism was heterozygous, displaying both the 129 and B6 alleles, indicating segmental loss of 129 chr14 distal to the DSB. Segmental loss was observed in 84% of LOH clones sorted at day 4 for both Cas9 and I-SceI DSBs (**Fig. 2A**), congruent with the FISH results (**Fig. 1H**), but increased to 98% in clones sorted at day 10 (**Fig. 2A**). The remaining day 4-sorted LOH clones were inferred to have undergone loss of the entire 129 chr14, as only the B6 allele was maintained at *Fhit*. Only a single day 10 clone was found with whole chromosome loss, indicating that these events were more likely to be outcompeted in the heterogeneous population prior to the sort.

A subset of clones from the day 10 timepoint was analyzed by whole chromosome painting. Of 6 clones identified to have segmental LOH by PCR, 4 had terminal deletions and 2 had nonreciprocal translocations of the centromere-proximal region of chr14 to an unidentified chromosome (**Fig. 2B**). The one clone with predicted whole chromosome loss had an absent 129 chr14 with duplication of the unbroken B6 chr14, indicating copy neutral LOH, which may account for its representation in LOH clones at day 10.

RNA-seq was performed on 4 of these clones to determine the effect of LOH on the transcriptional profile. Three clones with segmental loss – two from terminal deletion and one from translocation – demonstrated a significant decrease in expression of chr14 genes distal to the DSB (**Fig. 2C and S3A**). Of the ∼200 expressed genes in this region, ∼60 showed substantially decreased expression (≥1.5 fold lower), and as expected transcripts from this region were derived only from the B6 chr14 (e.g., **Fig. S3B**). No other such regional decrease in gene expression was observed for any of the other chromosomes (**Fig. S3C**). By contrast, levels of chr14 transcripts were unaffected in the clone with loss of the 129 chr14 and duplication of the B6 chr14 (copy neutral LOH). Thus, cells can survive DSB-induced LOH despite perturbations to gene expression from a large segment of a chromosome.

### Telomere addition and translocation of sequences to DSBs

Segmental loss from a DSB arises primarily from terminal deletions and less frequently from non-reciprocal translocations. To interrogate these events, we asked how well the centromere-proximal DNA end of the DSB was maintained in clones. We attempted to amplify a polymorphism at an absent HindIII site using a reverse primer located ∼100 bp centromeric to the DSB site (**Fig. S4A,B**). The polymorphism from the 129 chr14 could be amplified in a few clones from the day 4 sort (Cas9, 2/40; I-SceI, 3/29) but a larger fraction of clones from the day 10 sort (I-SceI, 21/56; p=0.0058) (**Fig. S4B,C**). The greater prevalence in day 10 clones suggests that cells with well-preserved centromere-proximal DNA ends had a growth advantage in the population compared to cells that had more extensive deletions. By contrast, a distal polymorphism (EagI site) was not amplified in any of the 56 tested clones from the day 10 sort, consistent with deletions distal from the DSB (**Fig. S4B,C**).

Chromosomes with terminal deletions are expected to have telomeres added to their ends to maintain stability during clonal expansion. We attempted to amplify telomeric sequences at the DSB using a reverse primer consisting of telomere repeats and a forward primer 340 bp proximal to the DSB at the HindIII polymorphism (**Fig. 2D**). Of the 26 clones that had a well-preserved centromere-proximal DNA end, telomere sequences could be amplified in 5 of them. In 4 clones, a *de novo* telomere was added at a 4 bp microhomology with the telomeric repeat in the I-SceI overhang. Telomere addition in the fifth clone was more complex and included a small deletion and insertions of sequences. We further attempted to amplify telomeres in the remaining 99 clones that do not have well preserved DNA ends by using a forward primer located 201 bp further centromere-proximal from the DSB. Telomere addition was detected in one additional clone, in this case at another 4 bp microhomology with the telomeric repeat located 124 bp from the DSB. Telomere additions likely remained undetected in other clones because the deletions extended beyond the binding site of the primer for amplification and/or there were large insertions at the DNA ends that precluded amplification.

To identify sequences that may be translocated to the DSB site, we performed inverse PCR on the 21 clones which maintained the HindIII amplicon but did not have detectable telomere addition (**Fig. S4D**).

Translocated sequences could be identified in seven clones, each of which were joined to the 129 chr14 DNA end with minimal processing (1 to 33 bp deletion). Three clones had translocated sequences from distinct genomic sites at chr15qA2, chr15qF3 (at a LINE1 element), and chr18qA2. Two clones had repetitive elements at the DSB end, a LINE1 element and a minor satellite repeat, that made it difficult to assign a genomic location. The remaining two clones had sequences from the transfected I-SceI expression vector at the DSB end, which has been observed previously^34^. None of the clones had breakpoint junctions mapping to previously identified cryptic I-SceI sites in the mouse genome^35^.

### LOH is also frequent after a DSB on chr6

Thus far, we have shown that a DSB can lead to extensive LOH on chr14 in mESCs. To test if LOH would be observed at other chromosomes, we integrated the fluorescent markers at the *Rosa26* locus on chr6 such that mRuby3 and dClover2 expression is driven by the endogenous promoter of the *Rosa26* lncRNA (**Fig. S5A-C**). A DSB was introduced on the B6 chr6 26 Mb centromere-proximal to *Rosa26* at position 86.8 Mb (**Fig. 3A**). LOH was readily observed in ∼8% of the population irrespective of whether *dClover2* or *mRuby3* was integrated on the B6 chr6 in these clones. Thus, extensive LOH after a DSB is not restricted to chr14.

**Figure 3.**
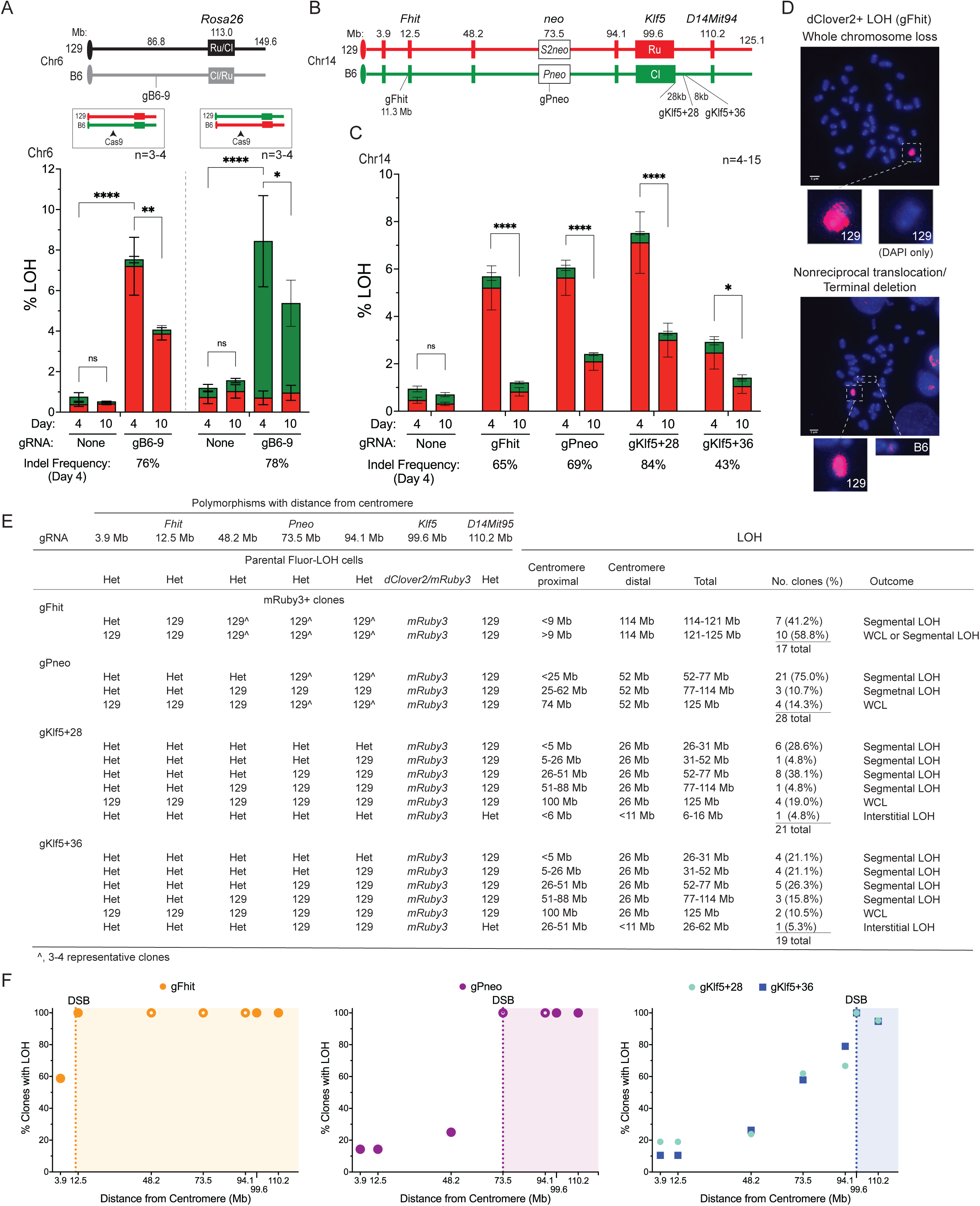
LOH can extend megabases centromere-proximal from the DSB. **A.** LOH is also observed using a Flo-LOH reporter integrated on chr6 at the *Rosa26* locus. A DSB was introduced 26 Mb centromere-proximal to *Rosa26* using a gRNA targeting B6 chr6, gB6-9. Two clones with reciprocal integration of *mRuby3* and *dClover2* were examined and found to undergo LOH at similar frequencies but in opposite directions. The percentage of cells that retain only dClover2 and mRuby3 are indicated by the green and red filled bars, respectively. Indel frequency was determined from the average of ICE analyses on the total cell population for all experiments. Error bars, mean ± s.d.; \*\*\*\**P*<0.0001, *\*\*P*=0.0038, *\*P*=0.0151 for total LOH; Ordinary one-way ANOVA, with Tukey’s multiple comparisons test. **B.** Four gRNAs were designed to specifically cleave the B6 chr14. The extent of LOH was determined using 7 polymorphisms located at indicated megabase distances from the centromere. The gRNAs that cleave distal to the *Klf5* gene, gKlf5+28 and gKlf5+36, cleave 28 and 36 kb, respectively, from the end of the *dClover2* expression cassette integrated at *Klf5*. **C.** LOH is induced by DSBs irrespective of chromosome position. Induced LOH is restricted to loss of dClover2 expression from the broken B6 chr14 and maintenance of mRuby3 expression from the unbroken 129 chr14. LOH is high in the population 4 days after Cas9 RNP transfection and declines significantly by 10 days in all cases. Indel frequency was determined from the average of ICE analyses on the total cell population. n=4-7 for each gRNA; n=15 with no gRNA. Error bars, mean ± s.d.; \*\*\*\**P*<0.0001, \**P*=0.0123 for total LOH; Ordinary one-way ANOVA, with Tukey’s multiple comparisons test. **D.** DSB induction in *Fhit* at 11.3 Mb from the centromere leads predominantly to LOH by whole chromosome loss. After DSB induction by Cas9, single-positive mRuby3 cell populations were sorted, and metaphase spreads were stained with DAPI and probed with a whole chr14 probe. While most spreads show loss of the entire B6 chr14 by whole chr14 painting, a portion have maintained a small fragment of the B6 chr14. Additional images in **Fig. S5A**. **E.** Extent of LOH after a DSB on the B6 chr14. Single-positive mRuby3 clones generated after sorting cells 4 days post-transfection were genotyped at the 7 polymorphisms. The inferred mechanisms of LOH are indicated. The extent of LOH for terminal deletions from the centromere-proximal DNA (left) end and centromere-distal (right) end are indicated, as is the total extent of LOH from both DSB ends. **F.** Summary of chr14 LOH after DSB formation from the four gRNAs. The percentage of clones with loss of the 129 polymorphism at each position is indicated. All clones had undergone LOH distal to the DSB (indicated by shading) except two from the *Klf5*-distal gRNAs which had undergone interstitial LOH. Filled circles indicate that all clones were genotyped at a polymorphism. Open circles indicate that only a subset of clones was genotyped typically because the indicated polymorphism was between polymorphisms that had fully undergone LOH or was at/distal to the DSB.

### Whole chromosome loss is common after a DSB near the centromere

To determine whether the chromosomal position of the DSB influences the frequency or type of LOH, we targeted a Cas9 DSB much closer to the centromere. The DSB was positioned 11.3 Mb from the centromere within the *Fhit* gene on the B6 chr14 (**Fig. 3B**). For comparison, a DSB was introduced 73.5 Mb from the centromere on the B6 chr14 at an integrated *neo* gene, at the same position of the previous 129 chr14 DSB. Four days after transfection, single-positive mRuby3 cells were observed at a similar frequency (∼6%) whether the DSB was targeted close to the centromere at *Fhit* or much more distal at *neo* (**Fig. 3C**). In both cases, LOH decreased between day 4 and day 10, implying that the cells with LOH in each condition were outcompeted by heterozygous cells, although the decrease was more pronounced with the DSB at *Fhit*.

However, chromosome painting of sorted mRuby3+ cells showed a major difference in outcome in that whole chromosome loss was much more frequent from the *Fhit* DSB (**Fig. 3D and S6A**). Segmental LOH from terminal deletions and nonreciprocal translocations was also observed in some cells with small regions of the B6 chr14 retained.

To characterize LOH events at greater resolution, we sorted individual single-positive mRuby3 cells four days after Cas9 transfection and analyzed chr14 markers. Clones derived from a DSB at either *Fhit* or *neo* exhibited LOH at all polymorphisms distal to the DSB site (**Fig. 3B,E)**. More than half of the *Fhit* clones also exhibited LOH at the most centromere-proximal polymorphism tested (at 3.9 Mb), as would be expected from whole chromosome loss, whereas many fewer clones with a DSB at *neo* exhibited LOH at this marker (**Fig. 3E and S6C**). The remaining *Fhit* clones maintained heterozygosity at the 3.9 Mb polymorphism, indicating segmental loss. Thus, while segmental loss is a frequent outcome from a DSB, a DSB closer to the centromere leads to more frequent whole chromosome loss.

### LOH frequently extends megabases centromere-proximal from the DSB site

To determine the potential for long interstitial deletions, we introduced a DSB distal to *Klf5*, reasoning that terminal deletion from the DSB site would not be sufficient to generate LOH. Targeting a Cas9 DSB either 28 or 36 kb distal to *Klf5* on the B6 chr14 (gRNAs gKlf5+28 and gKlf5+36, respectively) led to a large increase in single-positive mRuby3 cells, reaching >7% of cells for the more efficient gRNA, similar to a DSB at *Fhit* or *Pneo* (**Fig. 3B,C**). As expected, all 40 clones that were isolated from sorting at day 4 from the two gRNAs had lost the *dClover2* gene at *Klf5* (**Fig. 3E**). Surprisingly, only 2 clones maintained heterozygosity at the most distal polymorphism analyzed, *D14Mit95,* located 11 Mb from the DSB, which would be consistent with an interstitial deletion. Instead, most clones (32/40; 80%) were inferred to have undergone a terminal deletion, showing LOH at *D14Mit95* while maintaining heterozygosity at polymorphisms at 3.9 Mb and 12.5 Mb from the centromere.

To further characterize the extent of LOH, we examined several polymorphisms along the length of chr14 and were stunned to find that most clones had undergone LOH extending several megabases centromere-proximal to the DSB in addition to the distal LOH (**Fig. 3B,E and S6B-D**). LOH from the DSB towards the centromere was estimated to range from <5 Mb (10 clones), 5-26 Mb (5 clones), 26-51 Mb (13 clones), and 50-88 Mb (4 clones). Considering that the 8 clones in the most frequent size class from gKlf5+28 – a 26-51 Mb size range – comprise 38% of LOH events, and LOH occurs in ∼7% of the population, ∼2.5% of the total population have extensive loss. Thus, a DSB can generate huge swaths of LOH along the chromosome in both the centromere-proximal and distal directions. The remaining 6 clones had LOH at all polymorphisms examined, including the one located at 3.9 Mb, consistent with whole chromosome loss. This suggests that whole chromosome loss may be an extreme manifestation of the deletion process, rather than a distinct process.

A summary of LOH generated by the 4 gRNAs shows nearly universal LOH distal to the DSB with varying degrees of LOH centromeric to the DSB (**Fig. 3F**). As a proxy for cell fitness, we also compared the length of time for single cells to form colonies and expand (**Fig. S6E**). Despite loss of large amounts of genetic material, clones readily expanded in 20-30 days whether the DSB was located relatively close to the centromere (*Fhit*), near the middle of the chromosome (*Pneo*), or closer to the distal telomere (*Klf5*). However, the *Fhit* clones on average took longer to expand and had a wider range of expansion times, suggesting that whole chromosome or nearly whole chromosome loss may have a greater effect on fitness than segmental loss.

### Flo-LOH detects LOH arising from IH-HR which is increased with loss of BLM

DSB-induced recombination between chromosome homologs, termed interhomolog homologous recombination (IH-HR), leads to extensive LOH if it is associated with crossing over (**Fig. S7A**). Using an IH-HR reporter, we previously found that IH-HR is infrequent, and that the vast majority of IH-HR events are noncrossovers, such that extensive LOH from IH-HR is rare, making its detection laborious^32,36^. To facilitate the detection of LOH from IH-HR, the Flo-LOH system was introduced into cells containing the IH-HR reporter (**Fig. 4A**). This reporter consists of two nonfunctional *neo* genes integrated in the two chr14s, such that a DSB introduced into the *S2neo* gene (129 chr14) can be repaired from the *Pneo* gene (B6 chr14) to result in *neo*+ colonies. We found that a DSB from Cas9 or I-SceI induced IH-HR at similar frequencies (∼10^-4^) (**Fig. S7B**).

**Figure 4.**
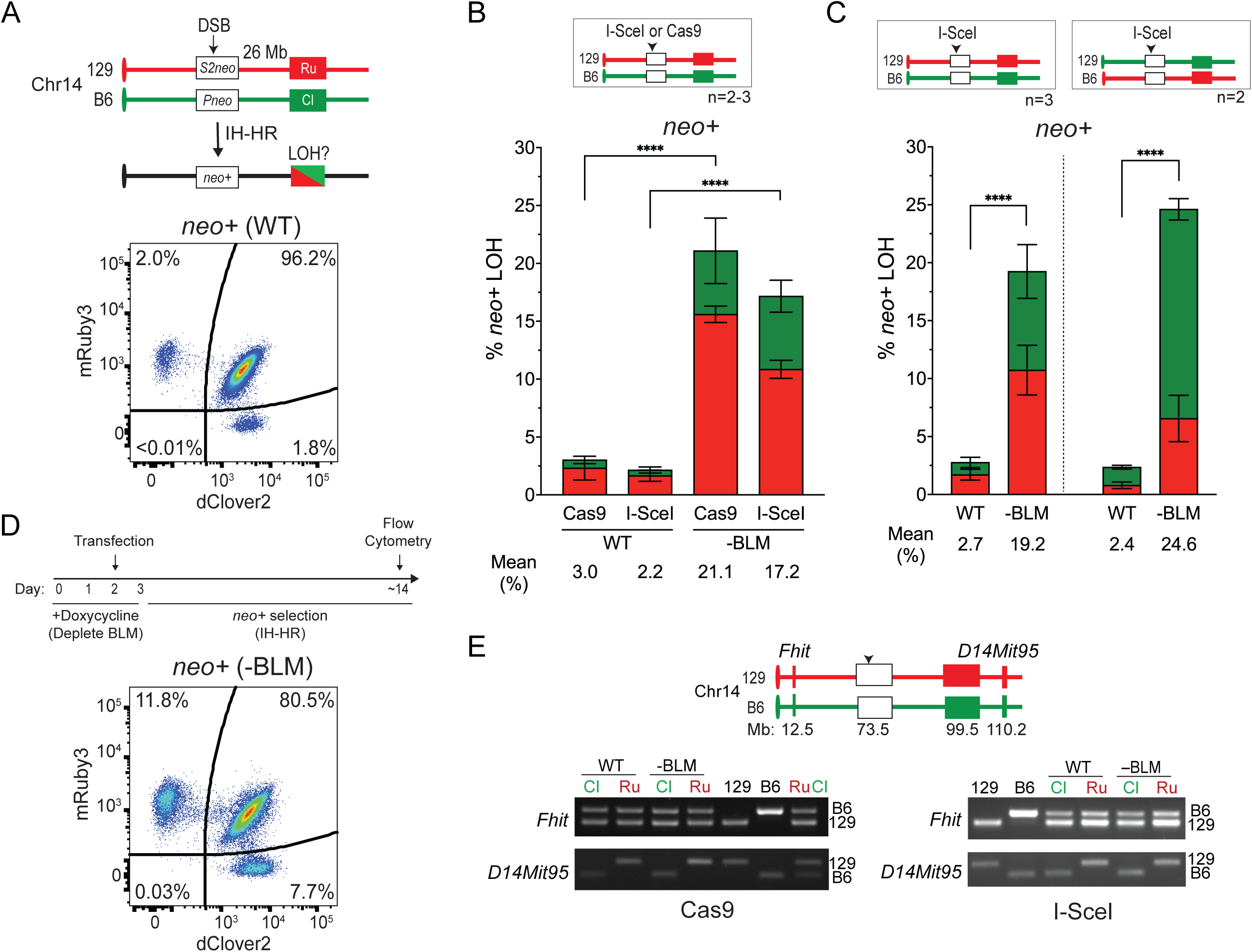
DSB-induced IH-HR leads to LOH which is suppressed by the BLM helicase. **A.** Interhomolog homologous recombination (IH-HR) reporter on chr14 in Flo-LOH cells. Two defective *neo* genes are integrated 26 Mb upstream of the fluorescent markers. IH-HR induced by a DSB leads to *neo+* cells and, if it involves crossing over, LOH from the DSB to the telomere. (See also **Fig. S6A**.) A DSB is required to generate *neo+* cells. Representative flow cytometric analysis following DSB induction and *neo+* selection shows an increase in single-positive mRuby3 and dClover2 cells compared to no DSB in unselected cells (Fig. 1B). **B,C.** DSB-induced IH-HR leads to LOH which is strongly suppressed by the BLM helicase. DSB induction by either I-SceI or Cas9 followed by selection for *neo+* cells leads to an increase in single-positive mRuby3 and dClover2 cells as compared with spontaneous LOH (Fig. 1B), which is further increased in the absence of BLM (**B**). Note that retention of only the 129 chr14 fluorescent marker is more common in both orientations than that of the B6 marker (**C**), as expected (**Fig. S6A**). n=2 clones for each orientation with 1-2 experiments per clone. Error bars, mean ± s.d.; \*\*\*\**P*<0.0001 for total LOH; Ordinary one-way ANOVA, with Tukey’s multiple comparisons test. **D.** Scheme of BLM depletion in IH-HR experiments. Representative flow cytometric analysis following BLM depletion, DSB induction, and *neo+* selection shows an increase in single-positive mRuby3 and dClover2 cells compared to wild-type cells without BLM depletion, as shown in **A**. **E.** Confirmation of LOH in single-positive *neo+* cell populations at the distal *D14Mit95* locus. By contrast, heterozygosity is maintained at the centromere-proximal *Fhit* locus, as expected from crossing over within the *neo* genes. RuCl, DNA from Flo-LOH 129/B6 parental cells; 129 and B6, DNA from 129 cells and B6 cells, respectively.

Flow cytometry of the *neo*+ populations demonstrated loss of one of the fluorescent markers in 2-4% of cells, indicative of LOH from crossing over (**Fig. 4A,B**). Similar results were obtained with both orientations of the fluorescent markers (**Fig. 4C**).

The BLM helicase, mutated in Bloom syndrome, is known for suppressing crossing over between sister chromatids as well as between homologs through its role in “dissolving” recombination intermediates that give rise to crossovers^32,37^. To test the effect of BLM loss in the Flo-LOH system, cells were treated with doxycycline prior to DSB formation to transiently suppress BLM expression from the modified endogenous alleles^33^ (**Fig. 4D**). In the absence of BLM, overall IH-HR occurred at a similar or slightly reduced frequency compared to wild-type cells^32,38^, but extensive LOH among *neo+* cells increased on average ∼8-fold (**Fig. 4B-D**), consistent with an increase in crossing over during IH-HR. In both wild-type and BLM-depleted cells, a skew was observed towards LOH events with the fluorescent marker from the 129 chr14, which is expected from the design of the recombination reporter (**Fig. S7A**). As would be expected for cells that experienced LOH during crossing over, wild-type and BLM-depleted *neo+* cell populations were heterozygous at the centromeric *Fhit* polymorphism but experienced LOH at the distal *D14Mit95* polymorphism (129 for mRuby3+ cells; B6 for dClover2+ cells) (**Fig. 4E**). Individual *neo+* clones gave similar results to cell pools, exhibiting LOH at the telomeric marker (**Fig. S7C**). Thus, Flo-LOH detects LOH from IH-HR and can be used to characterize factors that impact it.

### Genomic loss in the absence of BLM

Loss of BLM is associated with increased genomic instability as well as elevated crossovers. Therefore, we examined the effect of BLM depletion on LOH levels without selecting for IH-HR or introducing a DSB. Spontaneous LOH events were quantified by transiently treating cells with doxycycline for 3 days to suppress BLM expression and then performing flow cytometry 3 days later. Depletion of BLM led to a ∼6-fold increase in cells with LOH to nearly 4% of the population (**Fig. 5A**), indicating a high level of genomic instability in the population even without exogenous DNA damage. In this case, LOH was observed at a similar level for either fluorescent marker, as expected, since DNA damage was not specifically introduced on one of the chr14s.

**Figure 5.**
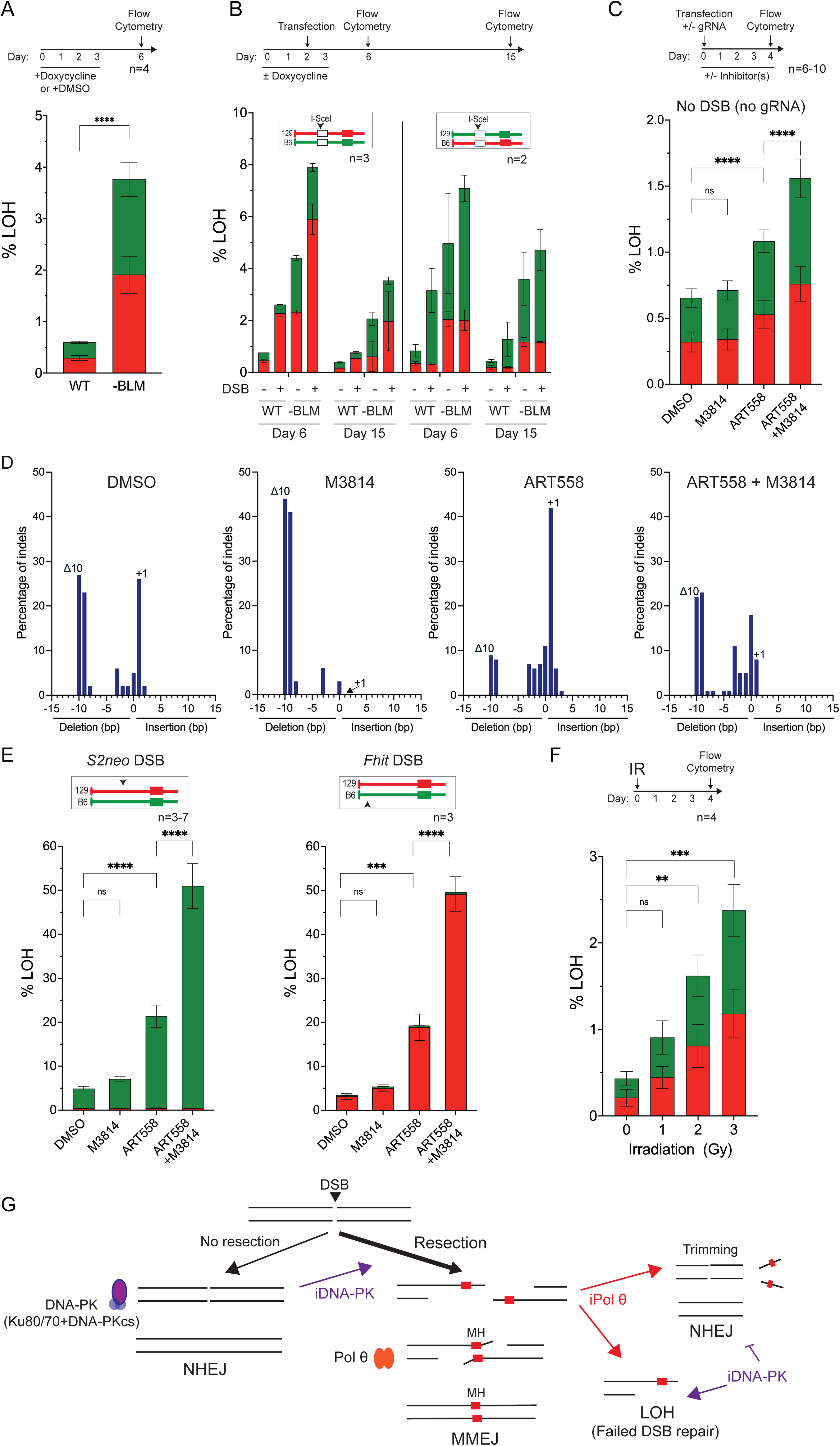
Inhibition of end-joining repair factors leads to a synergistic increase in LOH. **A.** LOH is enhanced by transient BLM depletion. Both single-positive populations are approximately equivalent after BLM depletion. 3 technical replicate plates in each experiment. Error bars, mean ± s.d.; \*\*\*\**P*<0.0001 for total LOH; Unpaired two-tailed t test. **B.** LOH is enhanced by transient BLM depletion but decreases over time. Note that with BLM depletion alone, both single-positive populations are approximately equivalent, while with a DSB, cells which have lost the marker on the broken chromosome are specifically increased in an additive manner. **C.** Inhibition of end joining increases spontaneous LOH. Although the impact of the DNA-PK inhibitor M3814 is minimal, Pol θ inhibition with ART558 significantly increases LOH, and combined inhibitor treatment leads to a further increase. Error bars, mean ± s.d.; \*\*\*\**P*<0.0001 for total LOH; Ordinary one-way ANOVA, with Tukey’s multiple comparisons test. **D.** Indel distribution after DSB repair in the presence of end joining inhibitors. The two main end joining products (+1 and Δ10) at the *S2neo* DSB are conversely affected by M3814 and ART588, consistent with inhibition of NHEJ and MMEJ, respectively. The Δ10 product occurs at a 2 bp microhomology, which is within a larger segment of imperfect microhomology (**Fig. S8B**). A Δ9 bp product at a 4 bp perfect homology also increases with M3814 and is suppressed by ART558, consistent with that product forming preferentially by MMEJ. Results are shown from one experiment; a summary of results from all experiments (n≥3) is in **Fig. S8C**. Sequences of end joining products are in **Fig. S8B**. **E.** DSB-induced LOH is sharply increased by inhibition of MMEJ (ART558) and synergistically increased by inhibition of both MMEJ and NHEJ (ART558+M3814) at both the *S2neo* (73.5 Mb) and at *Fhit* (11.3 Mb) loci. Error bars, mean ± s.d.; \*\*\*\**P*<0.0001, \*\*\**P*=0.0003 for total LOH; Ordinary one-way ANOVA, with Tukey’s multiple comparisons test. **F.** LOH is enhanced by irradiation-induced DNA damage in a dose-dependent manner. Both single-positive populations are similarly increased, as would be expected from random DNA damage to either chr14. Error bars, mean ± s.d.; \*\*\**P*=0.0001, \*\**P*=0.008 for total LOH; Ordinary one-way ANOVA, with Tukey’s multiple comparisons test. **G.** Interaction of the two end-joining pathways for LOH suppression in mESCs. NHEJ is not essential for suppressing LOH, because ends that fail to join by NHEJ due to DNA-PK inhibition can be resected and joined by MMEJ. However, when MMEJ fails due to Pol θ inhibition, resected DNA ends are inefficiently joined by NHEJ and may require trimming of resected overhangs. When both pathways are inhibited, LOH is frequent due to terminal deletions.

When a DSB was introduced into the 129 chr14, an additive increase in LOH specific to the broken chromosome was observed in the absence of BLM, reaching 7-8% of the cell population (**Fig. 5B**), suggesting that LOH from BLM depletion and DSB induction occurs via separate pathways. Cells with LOH decreased in the population over the next 9 days under both conditions, presumably due to a growth disadvantage. Thus, the impact of genetic perturbations on genomic integrity in the presence or absence of exogenous DNA damage are readily quantified using this system.

### Inhibition of both end joining pathways leads to a synergistic increase in LOH

To determine which biochemical pathways affect DSB-induced LOH, we investigated the two major end joining pathways in mammalian cells, canonical nonhomologous end joining (NHEJ) and microhomology-mediated end joining (MMEJ)^39^. NHEJ is considered to be the most common pathway to repair DSBs in mammalian cells. While it can lead to insertions and deletions at the DSB site, DNA ends are typically well preserved. MMEJ is a genetically separable pathway in which joining occurs at short microhomologies within resected DNA ends^40,41^. While traditionally considered to be a back-up pathway for DSB repair, for example, when HR fails^42,43^, recent work indicates it can operate without perturbation to other repair pathways, including at a Cas9 DSB^44,45^.

To examine the importance of the two end joining pathways in the prevention of LOH, we transiently inhibited factors within each pathway. The compound M3814 (peposertib) inhibits the catalytic activity of DNA-PK, impairing NHEJ^46^, whereas ART558 inhibits the DNA polymerase activity of Pol θ, hindering MMEJ^47^. In the absence of a Cas9-induced DSB, DNA-PK inhibition had no discernible effect on LOH, while Pol θ inhibition led to a 1.7-fold increase and combined DNA-PK and Pol θ inhibition led to a 2.4-fold elevation in LOH (**Fig. 5C and S8A**). Thus, MMEJ has an important role in suppressing LOH arising from endogenous lesions; in its absence, NHEJ also becomes important for suppressing LOH.

HR is available to repair endogenous DNA damage but is expected to be unavailable for Cas9-induced DNA damage if both sister chromatids undergo DSBs. Thus, LOH from Cas9-induced DSBs may have different dependencies than LOH arising from endogenous damage. We first tested the inhibitors for their impact on end joining and found that the inhibitors affected the expected pathways at a DSB either mid-chromosome at *S2neo* or nearer to the centromere at *Fhit*. The two main end-joining events at the *S2neo* DSB were a 1-bp insertion expected from NHEJ and a 10-bp deletion at a 2 bp microhomology expected from MMEJ (**Fig. 5D and S8B**). M3814 treatment eliminated the +1 bp product but augmented the Δ10 bp product, whereas ART558 treatment substantially diminished Δ10 bp product but increased the +1 bp product. A similar impact on two end joining products was observed at the *Fhit* DSB, although slightly less pronounced (**Fig. S9A,B**).

Thus, both NHEJ and MMEJ contribute to repair in unperturbed conditions at the DSBs, but the two pathways can backup each other when one is defective, at least to some extent. The complexity of the end joining profiles makes it difficult to ascertain whether one pathway can fully compensate for the other. However, considering the dominant end joining events, it appears that MMEJ can better compensate for loss of NHEJ than NHEJ can for loss of MMEJ (**Fig. S8C,S9C**).

Examining LOH, we found that DNA-PK inhibition led to a ∼1.6-fold increase in LOH whether the DSB was introduced at *S2neo* or *Fhit*, although this was not statistically significant (**Fig. 5E**). However, the impact on LOH was dramatic when Pol θ was inhibited: LOH increased ∼5-fold to ≥20% of cells with either gRNA. The combined inhibition of both Pol θ and DNA-PK led to a synergistic ∼12-fold increase in LOH, reaching an astonishing ∼50% of the population. Cell proliferation was not significantly affected by inhibitor treatment, except with the combined treatment which caused a decrease in cell number (**Fig. S8D,9D)**. These results demonstrate the importance of the two end joining pathways in the maintenance of genomic integrity upon DSB formation (**Fig. S8D,S9D)**, and moreover implicate MMEJ as the predominant DSB repair pathway to suppress LOH, even when NHEJ is functional.

### Genomic loss from global DNA damage from ionizing radiation

While Cas9 introduces a targeted DSB, treating cells with ionizing radiation leads to genome-wide DNA damage, including DSBs. For example, 1 Gy leads to ∼40 DSBs in the genome^48^, as well as numerous other lesions including single strand breaks. To test the impact of global DNA damage on LOH, cells were treated with sublethal doses of ionizing radiation. We found that irradiation led to a dose-dependent increase in LOH, up to ∼2% at 3 Gy (**Fig. 5F**). LOH was observed at a similar level for either chr14, consistent with a random distribution of DNA damage among the chromosomes. These results indicate that the Flo-LOH system can be used as a proxy for genome-wide instability arising from global DNA damage.

## Discussion

In this study, we established a two-color, flow cytometry-based approach to identify and quantify loss of heterozygosity (LOH) and interrogate factors that impact LOH. The Flo-LOH system was developed in mouse embryonic stem cells (mESCs) which have one set of chromosomes derived from B6 mice and one from 129 mice, such that a DSB can be introduced into a single chromosome homolog (B6 or 129 strain) and the extent of LOH mapped using existing polymorphisms between the two strains. With a focus on allele-specific DSB-induced events, we find that megabase-scale LOH occurs in a few percent of cells after a DSB, whether the DSB is centromere-proximal, distal, or more central within the chromosome. LOH entails large structural changes, e.g., terminal deletion, centromere-proximal deletion, nonreciprocal translocation, and whole chromosome loss. Ionizing radiation, which creates DNA damage genome wide, including DSBs, also induces significant levels of LOH. Cells with LOH are outcompeted when maintained within a population consisting mostly of heterozygous cells but can propagate when isolated from that population despite losing huge swaths of genetic information.

### Extensive LOH centromere-proximal from the DSB

When a DSB is targeted to the middle of the chromosome or nearer to the end of the chromosome, the majority of LOH events are segmental with loss of the distal part of the chromosome (52 Mb and 26 Mb, respectively). Terminal deletions can be explained by micronucleation of the acentric fragment resulting in its subsequent loss^23^. The centromeric chromosome fragment can be propagated and in fact is well preserved in a fraction of clones which have just a few bp deleted from the DNA end. However, in many other clones, the centromeric fragment exhibits extensive LOH proximal to the DSB site. With a DSB near the fluorescent markers, >40% of clones exhibit LOH ≥26 Mb from the DSB, and 10% have LOH extending ≥51 Mb. Since LOH at the makers is always continuous, these chromosomes with segmental loss do not appear to arise from a complex shattering and rejoining event like chromothripsis^23,49^. Resection without repair could lead to processive loss from DSB ends, but it is unlikely to reach megabase lengths, given that it is inherently slow. (Resecting 51 Mb at ∼4 kb/hr, the rate seen in yeast^50^, would take >10,000 hours.) We speculate that sister chromatids undergo DSBs contemporaneously and that the DNA ends fuse to form a dicentric chromosome. The dicentric chromosome then breaks when the cell attempts chromosome segregation, possibly through the action of the TREX1 nuclease^51^. If the DNA ends are not immediately protected, breakage-fusion-bridge cycles may continue^52,53^. Attrition of genetic material centromeric to the DSB has also been observed in human embryos, although on a scale of hundreds of kilobases instead of megabases^54^. This has been proposed to arise from protein-mediated tethering of sister chromatids at the DSB site causing secondary chromosome breakage after the chromatids begin to separate.

In addition to segmental LOH, we also observed whole chromosome loss leading to LOH. This was especially evident when a DSB is targeted relatively close to the centromere, which has also been observed in human embryos ^21^. It is possible that breakage close to the centromeric region prevents proper kinetochore interaction and spindle attachment, resulting in chromosome missegregation. Alternatively, given the extensive attrition that we observed accompanying many segmental LOH events, it is possible that whole chromosome loss events arise by the same attrition mechanism. To this end, a DSB near the centromere requires only ∼11 Mb loss from the proximal DNA end for the entire chromosome to be lost.

### Protection of DNA ends: telomere addition and translocation of sequences from other chromosomes

Chromosomes with segmental LOH require end protection to be stable. In some clones, chromosome ends were “capped” by another chromosome through a nonreciprocal translocation. Much more frequent were segmental LOH events with terminal deletions. In some clones that had well-preserved proximal DNA ends (∼≤100 bp loss from the DNA end), telomere sequences could be detected at or near the DNA ends at microhomologies with the telomere repeat. We expect these to arise from de novo telomere addition, since telomerase is active in mESCs and required for their long-term survival^55^ and de novo telomere addition at a Cas9 DSB has been reported in human cancer cells^56^. However, telomere addition could be observed in just a fraction of clones with well-preserved DNA ends (∼20%), possibly because additional sequences are present between the DNA end and the telomere which would suppress amplification. In clones with extensive loss from the proximal DNA end, direct telomere addition could not be determined by PCR due to the lack of highly resolved breakpoints. Alternatively, telomeres could be added through an alternative lengthening of telomeres (ALT)-like mechanism, although ALT is usually only observed in telomerase-deficient mESCs^57,58^. Interestingly, among the clones with segmental LOH analyzed for sequence additions to the proximal DNA ends, one had sequences derived from a subtelomeric chromosome band (15qF3, **Fig. S3D**). We anticipate the Flo-LOH system will be a valuable resource for understanding mechanisms of telomere addition to DNA ends.

### Flo-LOH: A general sensor for genomic instability

Flo-LOH is a fast, sensitive, and flexible dual-color method to measure LOH, and we foresee it having widespread applications. Because it is flow cytometry-based, tens of thousands of cells can be quickly and cost-effectively screened for LOH in either chromosomal direction. Although other approaches exist for measuring LOH, including FISH^20,22^, fluorescent porphyrin accumulation^29^, and sequencing methods^26,27^, the benefits of Flo-LOH include its accessibility and the ability to readily isolate cell populations or single cells with LOH at any chromosome by flow cytometry for further characterization. This is especially powerful because LOH outcomes after a DSB are so heterogenous and LOH cells compete with non-LOH cells. Additionally, this two-color system allows LOH to be quantified from either or both chromosomes after allele-specific DSBs. Flo- LOH is also temporally flexible, as one can interrogate increases in LOH beginning 24 hr after perturbation and continue as long as required to determine the impact of LOH events on cellular fitness.

In principle, Flo-LOH can be developed in any cell line that is genetically manipulable. We started with mESCs because they represent a key developmental stage and because of their many advantages, including a short cell cycle and easy clonability. Their derivation from F1 hybrid 129/B6 mice allows polymorphism analysis along the lengths of chromosomes and, moreover, the particular cell line we used has an integrated reporter allowing for selection of IH-HR events^32^ and also repressible *Blm* alleles^33^. Because LOH from IH-HR is infrequent, due to the suppression of crossing over, it is laborious to screen for such events by PCR^32^. By incorporating flow cytometry into the system, it is now feasible to screen for factors that impact LOH from IH-HR. Moreover, as exemplified by the effect of BLM depletion, Flo-LOH can also be used as a general sensor of genomic instability. In addition to genetic perturbations, genomic instability can be quantified after treatment with drugs and other small molecules. Finally, Flo-LOH can be used as a measure of large on-target genomic changes for other types of genome engineering, such as base editing or prime editing.

### Survival of cells after extensive LOH

Unexpectedly, we found that cells that undergo extensive LOH following a DSB can survive and proliferate. Although outcompeted when cultured with heterozygous cells, they have only minimally affected colony formation and proliferation rates when removed from the general population. Cells with well-preserved DNA ends (and thus less LOH) have less of a growth disadvantage compared with cells that have lost an entire chromosome. That cells with losses of tens of megabases of genomic information are reasonably fit is surprising given that mESC lines are notable for having a relatively stable genome^59,60^. When karyotype changes have been identified in mESC lines, many involve chromosome duplications or amplifications rather than deletions or other loss events^61–66^.

This raises the question as to whether out-competition of LOH cells is as common *in vivo* during embryonic development as it is *in vitro* where cells are rapidly proliferating. In a recent study, about half of Cas9-targeted human embryos experienced extensive LOH on the targeted chromosome^21^. Embryonic stem cell lines could not be developed from these embryos, indicating selection against genomically-compromised cells during development. Recent studies have suggested that p53 expression may suppress LOH in human T cells^27^ and immortalized foreskin fibroblasts^22,29^, and that decreased expression or knockout of p53 correlates with higher levels of LOH. mESCs do not undergo p53-dependent apoptosis or cell cycle arrest upon DNA damage^67^, which may contribute to the proliferation of cells with LOH that we observe.

### End joining inhibition leads to rampant DSB-induced LOH

Inhibition of the two end joining pathways dramatically impacts DSB-induced LOH. Although DNA-PK inhibition only marginally increases LOH, Pol θ inhibition results in extremely high LOH levels (20%), and combined inhibition leads to an astonishing ∼50% of cells with LOH. These results indicate that the NHEJ pathway is not essential for suppressing LOH when MMEJ is functional. However, when MMEJ is crippled, NHEJ becomes crucial. We hypothesize that upon NHEJ inhibition, LOH is largely prevented because DSBs that would normally be repaired by NHEJ are instead efficiently resected and shunted for repair by MMEJ (**Fig. 5G**). Because mESCs have a high S phase component^67^, they may be particularly primed to resect DNA ends. By contrast, when MMEJ is inhibited, LOH is high because resected DNA ends are poor substrates for NHEJ due to the reduced capacity of Ku to load onto 3’ overhangs^41^. Thus, when both pathways fail, LOH is extremely high.

Without a DSB, we observed a similar dependence on the two end joining pathways for preventing LOH as with a DSB: NHEJ inhibition has little effect, MMEJ inhibition has a significant effect, and inhibition of both pathways increases LOH synergistically. While the level of spontaneous chr14 LOH in the Flo-LOH assay is much lower than with a DSB directed to chr14, LOH is presumably elevated genome wide, such that the overall level of spontaneous LOH on 40 mouse chromosomes may be comparable to DSB-induced LOH specifically on one chromosome. Thus, endogenous DNA damage and Cas9-induced DSBs may have a similar reliance on the two end joining pathways.

Chemical inhibition of end joining pathways has been used to improve gene editing outcomes in several contexts. Dual inhibition of NHEJ and MMEJ has recently been reported to lead to increased HR with exogenous DNA substrates in mESCs and a variety of human cell lines^45^. In some cell lines inhibition of NHEJ alone had a larger effect on HR frequency than MMEJ inhibition, but notably MMEJ inhibition in induced pluripotent stem cells caused a larger increase in HR. Together with the large increase in LOH we observe with MMEJ inhibition, these results suggest that stem cells may be more dependent on MMEJ for DSB repair than differentiated cells. Identifying the on-target effects of Cas9 DSBs in stem cells is particularly important considering the recent clinical FDA approval of CRISPR-modified hematopoietic stem cells^68^. Modulating individual end joining pathways has also been proposed to promote desirable end joining products, for example, Pol θ inhibition to promote 1 bp insertions to restore the reading frame in truncating mutations^45,69^.

Our results suggest that such approaches need to proceed with caution and take into consideration whether LOH could confound such efforts, especially when using populations of cells where such events cannot be readily identified and removed.

### Limitations

ESCs are pluripotent and represent a key stage in embryo development and therefore are an important cell type to investigate. While mESCs have a high S phase component, which may impact LOH outcomes, human ESCs also experience a long S phase^70^. Nevertheless, it will be important to extend our studies to differentiated cell types in a variety of lineages. Additionally, although we have identified pathways that suppress LOH, how to harness these or other pathways to prevent LOH remains elusive. Flo-LOH allows for the easy identification and characterization of segmental or whole chromosome loss events. However, it is unable to track chromosome gains, which are also common in cancer cells, although methods have been developed to promote whole chromosome or arm-level gains through centromere dysfunction^71^.

## Acknowledgements

We thank members of the Jasin lab, especially Deepika Prasad, Matteo Ferrari, and Raymond Wang, for assistance and discussions. We are grateful for the expert technical support from the past and present members of the MSK core facilities, including the flow cytometry core facility, the integrated genomics operation, and the molecular cytogenetics core facility including Marta Lisi, Gouri Nanjangud, Kalyani Chadalavada, Prajakta Kokate, and Murray Tipping. This work was supported by T32HD060600 and 1F31CA268775 (S.B.R.), P30CA008748 and R35CA253174 (M.J.).

## Author contributions

S.B.R., D.M., and M.J. conceived the study; S.B.R. performed the experimental work with assistance from D.M., T.B.W., Y.-Z.J., S.J., and Q.D. and under the supervision of M.J.; S.B.R. and M.J. wrote the manuscript.

## Declaration of interests

The authors declare no competing interests.

**Supplemental information** Document S1. Figures S1-S9 Document S2. Tables S1-S3

## Methods Cell culture

The 129/B6 mouse embryonic stem cells (mESCs), containing one set of chromosomes from 129 mice and one set of chromosomes from B6 mice, were previously modified to contain a tet-repressible *Blm* promoter^33^ and an interhomolog homologous recombination (IH-HR) reporter^32^. These 129/B6 cells were further modified (as described below) to express mRuby3 and dClover2 from each allele of *Klf5* on chr14 or *Rosa26* on chr6 for detection of loss of heterozygosity. Cells were grown on gelatin-coated plates in Dulbecco’s Modified Eagle Medium (Memorial Sloan Kettering Cancer Center Media Preparation Core Facility) supplemented with 12.6% Stasis stem cell qualified FBS (GeminiBio #100-525), 1% Non-Essential Amino Acids (Gibco #11140-050), 1% L-glutamine (Memorial Sloan Kettering Cancer Center Media Preparation Core Facility), 1% Penicillin-Streptomycin (Gibco #15140-122), 0.008% murine leukemia inhibitory factor (GeminiBio #400-495), and 0.0007% beta-mercaptoethanol (Sigma #M7522). Cells were tested regularly for mycoplasma contamination using the Mycoplasma PCR Detection Kit (Applied Biological Materials #G238).

## Oligonucleotide sequences

Relevant information about all primers, gRNAs, and plasmids, as well as sequences for all primers and gRNAs, are listed in **Tables S1-S3**.

### Targeting *mRuby3* and *dClover2* to *Klf5* to generate Flo-LOH on chr14

Template plasmids for targeting *mRuby3* and *dClover2* to the C terminus of *Klf5* were created using NEBuilder Hifi DNA Assembly (New England BioLabs #E2621). The backbone, pBluescript SK+, was digested with EcoRV (New England BioLabs #R0195), and all fragments had 10-20 bp overlap with the adjacent fragment or backbone. The ∼500 bp fragments containing the homology arms were amplified from 129/B6 mESC genomic DNA using Q5 high-fidelity DNA polymerase (New England Biolabs #M0491).

The 5’ homology arm, containing the sequence of *Klf5* exon 4 and part of intron 3, was amplified from mESC genomic DNA using primers Klf5 ex4 5’ hom_fwd and Klf5 ex4 5’ hom_rev_T2A. Given that the amplified sequence contains the gRNA binding sequence and PAM, two alterations were introduced into the 5’ homology arm to be subsequently incorporated into the edited alleles. The sequence corresponding to the PAM site was disrupted with introduction of a G to C point mutation at the second G of the PAM site (TGG to TGC) to prevent Cas9 cutting of the edited allele. The *Klf5* stop codon TGA was deleted, with the dual purpose of allowing expression of the T2A-fused fluorescent markers and disrupting the gRNA sequence to further prevent Cas9 cutting of edited alleles. The 3’ homology arm, containing the 3’UTR and part of the intergenic region downstream of *Klf5*, was amplified using primers Klf5 ex4 3’ hom_fwd_Ruby or Klf5 ex4 3’ hom_fwd_Clover and Klf5 ex4 3’ hom_rev.

The *mRuby3* fragment was amplified from a mRuby3 expression plasmid derived from pKanCMV- mClover3-mRuby3^72^ (Addgene #74252) using Q5 high-fidelity DNA polymerase (New England BioLabs #M0491) and primers T2A+mRuby3_for and mRuby3_rev to add the T2A sequence to mRuby3. The *dClover2* fragment was amplified from the dClover2-C1 plasmid^72^ (Addgene #54577) using Q5 high-fidelity DNA polymerase and primers T2A+dClover2_for and dClover2_rev to add the T2A sequence to dClover2. The T2A and fluorescent marker gene fragments were then PCR purified and amplified with the T2A_fwd and mRuby3_rev or dClover2 _rev primers for cloning. The Cas9 gRNA targeting *Klf5* (gKlf5) was cloned into the pSpCas9(BB)-2A-Puro plasmid^73^ (pX459, Addgene #48139) at the BbsI (New England BioLabs #R3539) restriction enzyme site. All minipreps were performed with PureLink Quck Plasmid Miniprep Kit (Invitrogen #K2100-03). All maxipreps were performed with the PureLink HiPure Plasmid Filter Maxiprep Kit (Thermo Fisher Scientific #K2100-17). All primers for this cloning, PCR, and sequencing were ordered from Integrated DNA Technologies.

To target *mRuby3* and *dClover2* to *Klf5*, 3 x 10^6^ modified 129/B6 cells were electroporated with 20 μg of each of the gKlf5 plasmid, the Klf5-dClover2 template plasmid, and the Klf5-mRuby3 template plasmid. Cells were electroporated in 650 μl opti-MEM reduced serum media (Gibco #11058-021) with a Gene Pulser Xcell (Bio-Rad) in a 0.4 cm cuvette (Bio-rad #1652091) with 250 V voltage, 950 µF capacitance, and infinite resistance. Seven days later, cells double-positive for mRuby3 and dClover2 expression were single cell sorted by flow cytometry on a FACSAria cell sorter (BD Biosciences) by the Memorial Sloan Kettering Cancer Center Flow Cytometry Core Facility to generate clones.

Double-positive mRuby3 and dClover2 expression was confirmed by flow cytometry of individual clones. Correct integration of *mRuby3* and *dClover2* at *Klf5* was confirmed using DreamTaq Master Mix (Thermo Fisher Scientific #K1082) PCR. Out-Out PCR was performed using primers Klf5 ex4 nesting_for and Klf5 ex4seqR3. Correct *mRuby3* integration at *Klf5* was confirmed by Out-In PCR with primers Klf5 ex4 nesting_for and Ruby int R1 and In-Out PCR with primers Ruby int F1 and Klf5 ex4 nesting_rev. Correct *dClover2* integration at *Klf5* was confirmed by Out-In PCR with primers Klf5 ex4seqF3 and dClover out in R1 and In-Out PCR with primers Clover seq F and Klf5 ex4 nesting_rev. Clones with correct integration of *mRuby3* and *dClover2* at *Klf5* were termed Flo-LOH clones.

### Targeting *mRuby3* and *dClover2* to *Rosa26* to generate Flo-LOH on chr6

Template plasmids for targeting *mRuby3* and *dClover2* to the *Rosa26* locus were created using a *Rosa26*- targeting plasmid^74^ (p15a_Kan_MLS_Rosa26_Short_CRISPRDel; Addgene #97008) and assembled using NEBuilder Hifi DNA Assembly. The *Rosa26* donor DNA was digested with PacI (New England BioLabs #R0547), and all fragments had 10-20 bp overlap with the adjacent fragment or backbone. The *mRuby3* fragment was amplified from the pKanCMV-mClover3-mRuby3 using Q5 high-fidelity DNA polymerase and primers mRuby3-SA-Koz-F and PR Ruby R. The *dClover2* fragment was amplified from the dClover2-C1 plasmid using Q5 high-fidelity DNA polymerase and primers dClover2-SA-Koz-F and PR Clover R. The SV-40 polyA signal sequence was appended to both *dClover2* and *mRuby3* by amplifying it from dClover2-C1 plasmid using primers PR SV40 F and dClover2-SV40pA-PacI-R and assembling it in the HiFi reaction.

To target *mRuby3* and *dClover2* to *Rosa26*, 2 x 10^6^ modified 129/B6 cells were transfected with 20 μg of each of the Rosa26-mRuby3 template plasmid, the Rosa26-dClover2 template plasmid, and the px330_Rosa26_sgRNA vector^74^ (Addgene #97007). Nucleofections were performed in 100 μl Homemade (HM) Amaxa-compatible buffer^75^ in 0.2 cm cuvettes (Bio-rad #1652086) using program A-013 of the Nucleofector 2b (Lonza). Thirteen days later, cells double-positive for mRuby3 and dClover2 expression were single cell sorted by flow cytometry on a SH800 cell sorter (Sony Biotechnology) to generate clones.

Double-positive mRuby3 and dClover2 expression was confirmed by flow cytometry of individual clones. Correct integration of *mRuby3* and *dClover2* at *Rosa26* was confirmed using Q5 high-fidelity DNA polymerase for 5’ Out-In and DreamTaq Master Mix for 3’ In-Out PCR. Correct *mRuby3* integration at *Rosa26* was confirmed by Out-In PCR with primers Ray-Rosa26-5’OutF1 and Ruby int R1 and In-Out PCR with primers Ruby int F1 and Ray-Rosa26-3’OutR2. Correct *dClover2* integration at *Rosa26* was confirmed by Out-In PCR with primers Ray-Rosa26-5’OutF1 and dClover out in R1 and In-Out PCR with primers Clover seq F and Ray-Rosa26-3’OutR2. Clones with correct integration of *mRuby3* and *dClover2* at *Rosa26* were termed *Rosa26* Flo-LOH clones.

## Transfection of Flo-LOH clones for LOH analysis

To induce a DSB using Cas9 RNPs, 2 x 10^6^ Flo-LOH cells were transfected with 140 pmol trRNA, 140 pmol crRNA, and 120 pmol Cas9. All trRNAs and crRNAs (Table) and were ordered from Integrated DNA Technologies; *S. pyogenes* Cas9 protein with an NLS was obtained from QB3 MacroLab (UC Berkeley).

Nucleofections were performed in 100 μl HM buffer in 0.2 cm cuvettes using program A-013 of the Nucleofector 2b. Following transfection, ∼2 x 10^5^ cells were generally plated on a gelatinized 10-cm plate. Cells were analyzed four days after transfection and, where relevant, passaged until analysis 10 days after transfection. For all flow cytometry, 50,000 DAPI negative (live) (Invitrogen, #D21490) cells per sample were read on the LSR Fortessa analyzer (BD Biosciences).

To induce a DSB at the I-SceI site at the *S2neo* gene, located at position 73.5 Mb on the 129 chr14, Flo-LOH cells were transfected with plasmids expressing I-SceI^76^ (pCBASceI, Addgene #26477) or Cas9- gRNA. The Cas9-gRNA expression plasmid, termed pX459+gS2Neo, was derived from cloning the gS2Neo into the pSpCas9(BB)-2A-Puro plasmid (pX459) at the BbsI restriction enzyme site. Transfection of 4.5 x 10^6^ cells was accomplished using 20 μg plasmid in 100 μl HM buffer in 0.2 cm cuvettes with program A-013 of the Nucleofector 2b.

Each *Klf5* Flo-LOH time-course involved three plus DSB transfections and one minus DSB transfection in parallel for each condition and was performed twice. Cells were plated after transfection in varying dilutions and then were analyzed at indicated time points or passaged for later time points.

For end joining inhibitor experiments, 2 x 10^5^ cells were plated in a gelatinized 10-cm dish after transfection and treated with inhibitors at the indicated concentrations. Cells were harvested 4 days post-transfection for flow cytometry analysis. ART558 (#HY-141520) was purchased from Cedarlanelabs, and M3814 (#S8586) was purchased from Selleckchem. Chemicals were dissolved in DMSO and stored in small aliquots at -80°C.

## Estimating cutting efficiencies

Cas9 cutting efficiencies with various gRNAs were estimated by quantifying indel frequency using sequence trace decomposition analysis with the ICE (Synthego) online software. Genomic DNA was isolated 3 days (time course experiments) and 4 days (Fig. 3) after transfection using the Puregene Core Kit A (Qiagen #158267). Amplicons of ∼1 kb surrounding the DSB site were derived using Dreamtaq polymerase (Thermo Fisher Scientific #EP0702) or Dreamtaq Master Mix, purified with Purelink Quick PCR Purification Kit (Invitrogen #K3100-02), and Sanger sequenced with a primer binding ∼200 bp from the DSB site. The gRNA, primer pairs, and sequencing primers are as follows: gS2neo: Primers e+h with Primer g; gChr6-B6-DSB9: Primers A-A9 B6 F2 + A-A9 B6 R2 with Primer A-A9 B6 F1 (experiments 1 and 2) or A-A9 B6 R1 (experiment 3); gPneo: Primers f+h with Primer g; gFhit: F-B6 gFhit and R-B6 gFhit with B6 gFhit ; gKlf5+28: F1-B6 gDwnstr Klf5+28 and R-B6 gDwnstr Klf5+28 with B6 gDwnstr Klf5+28 ; gKlf5+36: F2-B6 gDwnstr Klf5+36 and R-B6 gDwnstr Klf5+36 with B6 gDwnstr Klf5+36 seq.

Cutting efficiencies at the I-SceI site comparing I-SceI DSBs to Cas9 DSBs used a T7 endonuclease assay (New England BioLabs #M0302). Cells were transfected with plasmids expressing I-SceI (pCBASceI) or Cas9 (pX459+gS2Neo). Three days after transfection, genomic DNA was extracted using the Puregene Core Kit A. The region around the DSB site was amplified with Primers e+h with Dreamtaq polymerase. The PCR reaction was mixed 1:1 with 2x New England BioLabs Buffer 2, heated to 95°C for 5 minutes, decreased to 25°C by 0.1°C/second, cooled to 4°C for at least 15 minutes. The re-annealed PCR product was digested with 1.5U T7 endonuclease for 20 minutes.

## Colony formation and proliferation assay

Flo-LOH cells were transfected with plasmids expressing I-SceI or Cas9 and gS2neo to cleave the I-SceI site. Four days later dClover2+ cells, heterozygous cells, and all DAPI negative (live) cells were sorted via flow cytometry with an SH800 cell sorter (Sony Biotechnology). For each colony formation analysis, 1,000 and 2,000 cells were plated into 10-cm plates. Cells were grown until colonies were clearly visible, usually around 10-11 days. Colonies were fixed in methanol, stained with 2.7% Giemsa (Harleco, MilliporeSigma #620G-75) diluted in distilled water, and counted using the OpenCFU software^77^. The experiment was performed 2-4 times for each 1,000 and 2,000 cells plated for both I-SceI and Cas9. The graphed colony formation data shows the number of colonies normalized to the “all” cells condition for each quantity. Each point is the average of the two normalized values for that experiment. To confirm cell purity, plates from one experiment were analyzed with flow cytometry. Plates from “all” cells and heterozygous cells were ≥99.0% positive for both mRuby3 and dClover2 expression; plates from dClover2+ cells were ≥98.8% positive for dClover2 expression only.

For proliferation assays, cells were transfected with a Cas9+gS2neo RNP and sorted four days later (SH800 cell sorter). For each condition, 3-4 sorted populations of 50,000 cells each were plated into individual wells of a 12-well plate. Cells were trypsinized and counted every 48 hours, and 50,000 cells were plated again. This experiment was performed three times from independent transfections. On the last time point, cells were analyzed via flow cytometry to confirm purity of the sort. “All” cells and heterozygous cells were ≥95.7% and ≥98.9% positive for both mRuby3 and dClover2, respectively; wells from dClover2+ cells were ≥95.9% positive for dClover2 expression only.

## Whole chromosome painting

Cell populations were sorted by flow cytometry four days after transfection of the I-SceI expression plasmid or Cas9 RNPs, and harvested three days after sorting. Whole chromosome painting was performed using protocols established by the Memorial Sloan Kettering Cancer Center Molecular Cytogenetics Core Facility and a red mouse chr14 whole chromosome paint (Applied Spectral Imaging #FPRPR0169).

## Genomic analysis for LOH

Cells expressing a single fluorescent marker after a DSB were sorted 4 or 10 days after transfection into a 96- well plate with the SH800 or FACSAria cell sorters. Clones were expanded and confirmed by flow cytometry to be uniformly single-positive for dClover2 or mRuby3 expression (224 total), except for 8 which showed some level of heterozygosity (6 from gKlf5+28; 2 from gKlf5+36) and were subsequently discarded. Genomic DNA was extracted from LOH clones using the Puregene Core Kit A for further analysis with PCR. A few additional clones were then discarded because they maintained the gene for the absent fluorescent marker (3 clones, 1 with a mutation in *mRuby3* and 1 with a mutation in *dClover2*) or had not maintained the fluorescent marker gene but did not show LOH at or near the DSB site (2 clones).

PCRs for LOH analysis were done using Dreamtaq Master Mix and products were purified using the Purelink Quick PCR Purification Kit. Genotyping at the chr14 3.9 Mb polymorphisms was performed using allele-specific primers (Ube2e2_unv_F, Ube2e2_129_R, Ube2e2_B6_F, and Ube2e2_B6_R) together in one reaction to amplify different-sized products for each allele. Genotyping at *Fhit* (chr14 12.5 Mb) polymorphisms was performed using allele-specific primers (Fhit 129_3_3SNPs as F1, Fhit 129_3_3SNPs as R1, Fhit B6_2SNPs as F1, and Fhit B6_2SNPs as R1) together in one reaction to amplify different-sized products for each allele. Genotyping at the chr14 48.2 Mb SNP was performed by PCR amplification with non-allele-specific primers (48.2 For and 48.2 Rev)^32^, followed by Sanger sequencing (Azenta Life Sciences and Eton Bioscience).

Genotyping at the chr14 73.5 Mb-integrated mutant *neo* genes, specifically at the I-SceI and NcoI sites, was performed by PCR amplification using non-allele-specific Primers g+h^32^ and digestion with I-SceI (Thermo Fisher Scientific #ER1771) or NcoI (New England BioLabs #R0193). The B6-specific HindIII restriction site was genotyped by non-allele-specific PCR amplification with Primers a+b^32^ and digestion with HindIII (New England BioLabs #R0104). The 129-specific EagI and B6-specific PacI restriction sites were genotyped by non-allele-specific PCR amplification using Primers c+d^36^ and digestion with PacI or EagI-HF (New England BioLabs #R3505). Further allele-specific analysis around the DSB site was performed with universal Primer d and either 129-specific Primer e or B6-specific Primer f. PCRs to amplify sequences from de novo telomere addition were performed using 3 Telomere Repeats and either Primer a or Primer e.

Genotyping at chr14 94.1 SNP was performed by PCR amplification with non-allele-specific primers (94.1 For and 94.1 Rev)^32^ and Sanger sequencing. Amplification of *dClover2* and *mRuby3* used primers T2A_fwd and dClover2_rev or mRuby3_rev, respectively. Genotyping at the chr14 110.2 Mb (TG)_n_ polymorphism was performed by using non-allele-specific primers (D14Mit95 For and D14Mit95 Rev)^78^.

## RNA-seq

RNA sequencing was performed on one wild-type Flo-LOH sample and four LOH clones by the Memorial Sloan Kettering Cancer Center Integrated Genomics Operation Core Facility using the TruSeq stranded mRNA (polyA) library (Illumina) with a coverage of 20-30 million reads per sample.

## Inverse PCR

Inverse PCR was performed according to previously published methodology^79^. Restriction enzymes PsiI-v2 (New England BioLabs #R0744), AflII (New England BioLabs #R0520), ScaI (New England BioLabs #R3122), and NsiI (New England BioLabs #R0127) were chosen for analysis because they have only one recognition site each within 850 bp proximal to the I-SceI site. 5 μg of genomic DNA was digested in a 100 μl reaction overnight with 1 μl enzyme spiked in for the last hour of the digest. Digested DNA was purified using the PureLink Quick PCR Purification Kit, ligated with T4 DNA ligase (New England BioLabs #M0202), and then purified with phenol-chloroform (VWR #97064-694). Nested PCR was performed on the ligated DNA using Platinum II Taq Hot-Start DNA Polymerase (Invitrogen #14966005). The first round used Primer a and iPCR primer 9, and the second round used iPCR primer 8 and Primer e. This final nested PCR was purified using the PureLink Quick PCR Purification Kit or gel extracted using the PureLink Quick Gel Extraction Kit (Invitrogen #K2100-12) followed by Sanger sequencing.

## Selection for interhomolog recombination

For experiments selecting for interhomolog recombination (IH-HR) events, cells were nucleofected with I-SceI or Cas9 and gS2neo expressing plasmids. Media was removed 24 hours after transfection and replaced with fresh media containing 200 µg/ml G418 (GeminiBio #400-111P) to select for *neo+* cells. The G418-containing media was replaced every 3-4 days until all cells on the “no DSB” control transfection plates were dead and observable colonies formed on the DSB-induced plates (9-14 days). For experiments in which BLM was depleted, cells were treated with 1 µg/ml doxycycline 48 hours before transfection and plated into 1 µg/ml doxycycline directly after transfection. Media was replaced with doxycycline-free media 24 hours after transfection. Dox was dissolved in DMSO and stored in small aliquots at -80°C.

Electroporation was used in the experiments shown in Figure 4C and 4F. 15 x 10^6^ cells were electroporated with 60 µg I-SceI plasmid in opti-MEM reduced serum media (Gibco) with a Gene Pulser Xcell (Bio-Rad) in a 0.4 cm cuvette at 250 V, 950 µF, and infinite resistance. 3 x 10^6^ cells were plated in each 10-cm plate. All other preparations and treatment of the protocol excluding these transfection steps are identical to that of nucleofected cells.

## BLM depletion

To determine the effect of BLM depletion on LOH, 1-5 x 10^5^ cells were plated in a 6-cm plate. The following day, cells were treated with 1 µg/ml doxycycline (Millipore Sigma #D9891), which was maintained for 3 days. Doxycycline was removed on the third day and replaced with fresh media, and LOH was analyzed by flow cytometry three days after removal of doxycycline.

## Ionizing Radiation

To determine the effect of ionizing radiation on LOH, 1-10 x 10^4^ cells were plated in a 6-cm plate. The following day, cells were treated with 0-3 Gy of radiation using the X-Rad225 irradiator (Precision X-Ray Inc). LOH was analyzed by flow cytometry 4 days after irradiation.

---

**Figure S1.**
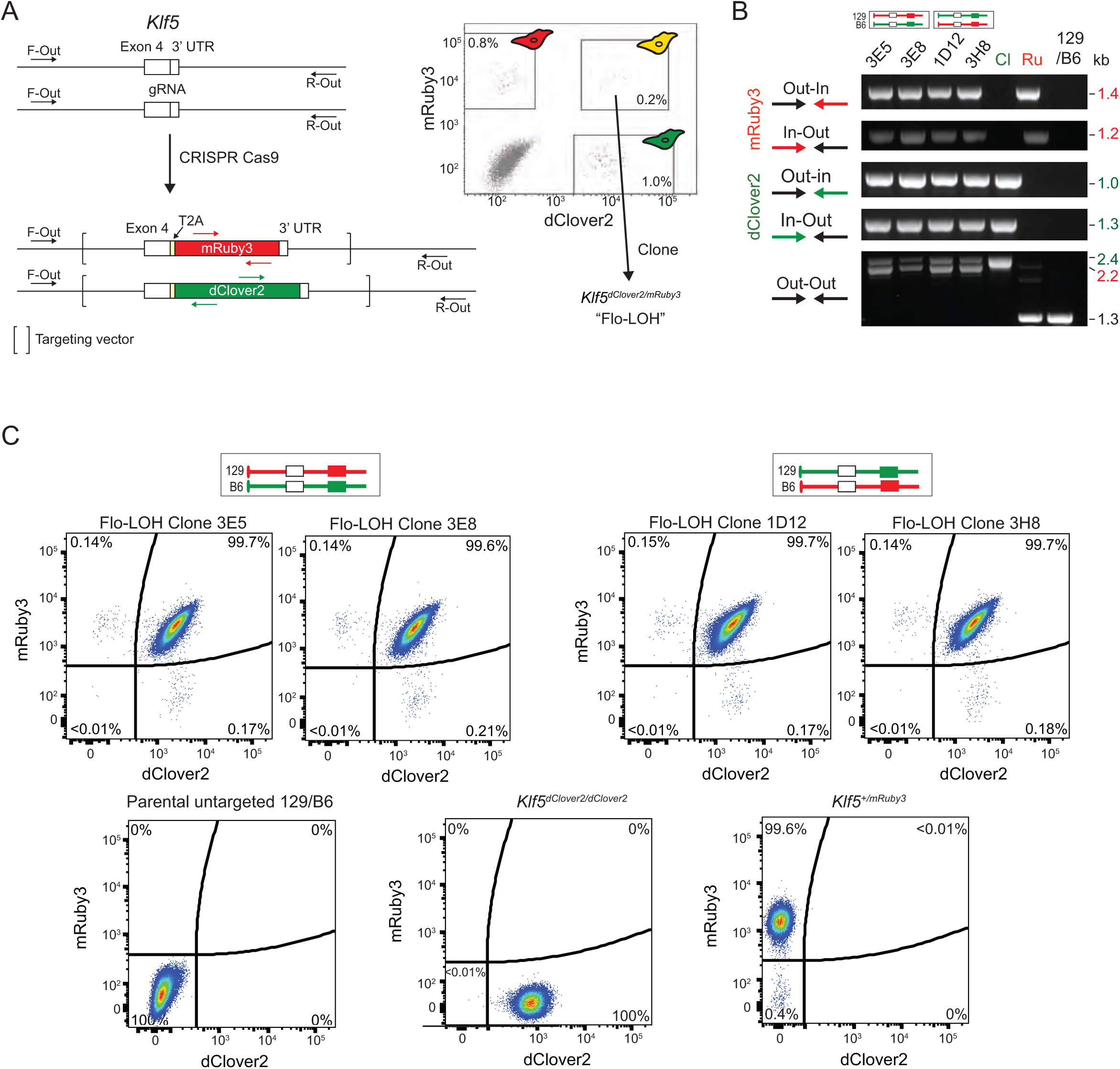
Flo-LOH: Targeting dClover2 and mRuby3 to the *Klf5* locus in 129/B6 mouse embryonic stem cells (mESCs). **A.** mESCs were transfected with a Cas9 and gRNA expression vector and *Klf5* targeting vectors with *mRuby3* and *dClover2* coding sequences. The fluorescent markers are promoterless but become expressed upon correct fusion to exon 4 at the 3’ end of *Klf5* through a T2A peptide sequence. The brackets delineate the extent of the *Klf5* homology arms in the targeting vectors. Double-positive cells were sorted from the transfected population to single cells by flow cytometry to generate Flo-LOH cells (*Klf5^dClover2/mRuby3^*). Single-positive cells were also sorted to generate compensation controls for subsequent flow cytometry. F-Out and R- Out represent the location of forward (Klf5 ex4 nesting_for and Klf5ex4seqF3) and reverse (Klft ex4 nesting_rev and Klf5 ex4seqR3) primers, respectively. **B.** PCR confirmation of correct dual integration at the *Klf5* locus. Primer locations are shown in **A**. Out-out primers are upstream and downstream of the homology arms in the *Klf5* locus. Correctly targeted *dClover2* and *mRuby3* insertion gives rise to ∼2.4 and 2.2 kb amplicons, respectively. Out-in and in-out primers use one of the out-out primers and another primer within the fluorescent marker, as indicated. Both orientations were recovered: clones 3E5 and 3E8 have integrations of *mRuby3* on the chr14 derived from the 129 mouse strain and *dClover2* on the chr14 derived from the B6 strain (*mRuby3*^1^^29^*;dClover2*^B6^), while clones 1D12 and 3H8 have the opposite orientation (*dClover2*^129^;*mRuby3*^B6^). The *Klf5^dClover2^*clone shown here (Cl) has integration of *dClover2* into both *Klf5* alleles. The *Klf5^mRuby3^* clone (Ru) has integration of *mRuby3* into a single *Klf5* allele, although in the out-out PCR the smaller band from the untargeted allele amplifies much better than the targeted allele. 129/B6, parental F1 hybrid cells. **C.** Flo-LOH clones in both orientations uniformly express dClover2 and mRuby3, as seen by flow cytometry, with minimal spontaneous LOH (0.3-0.4%). Cells expressing only dClover2 or mRuby3 were used for controls in PCR and flow cytometry experiments.

**Figure S2.**
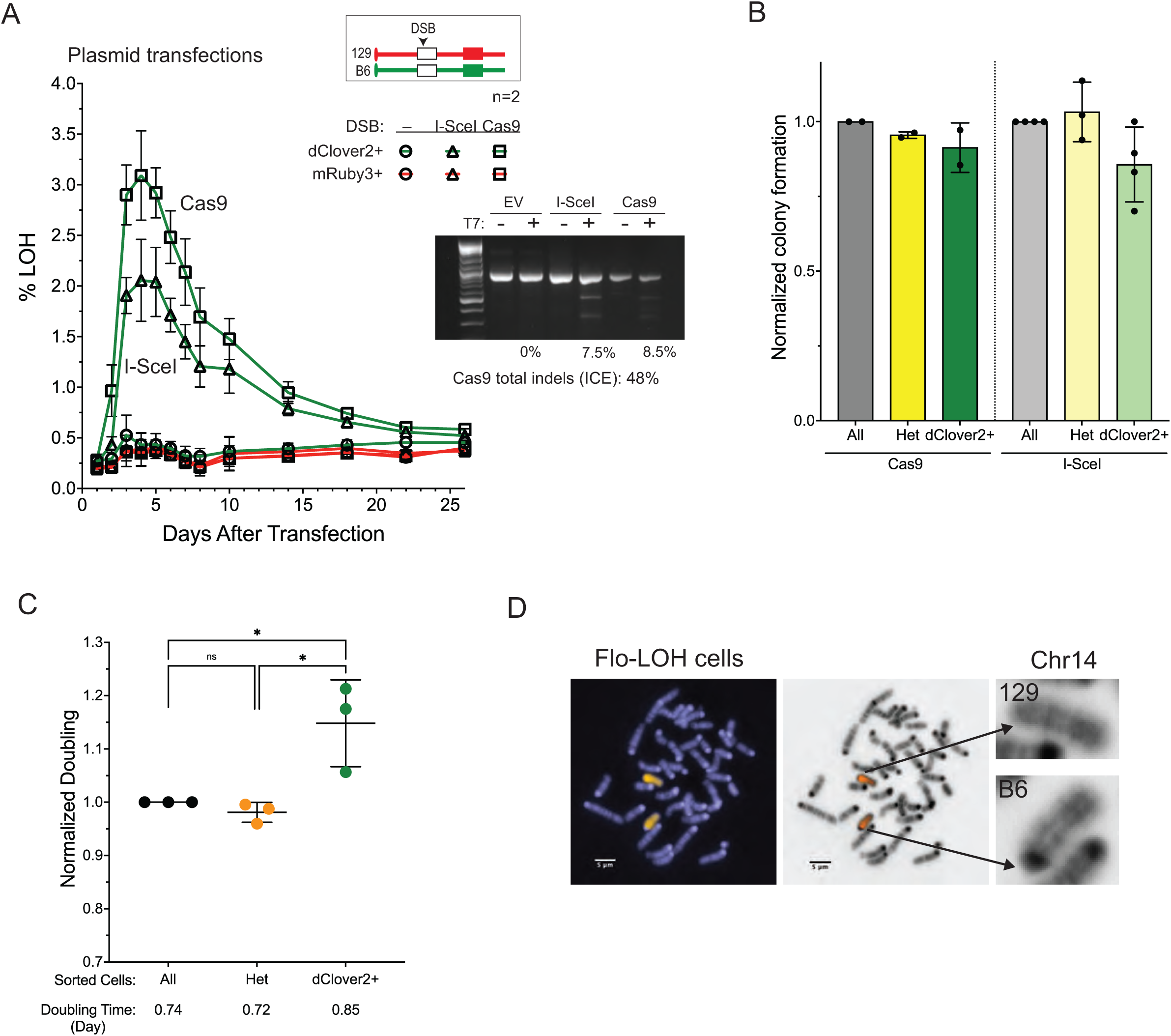
LOH is transiently observed in a population, but LOH cells are viable and proliferate. **A.** DSB-induced LOH increases then decreases over time. DSBs were introduced after transfection of plasmids expressing I-SceI or Cas9. The percentage of cells with LOH in the population peaks at a few percent after ∼4 days, then decreases over time to nearly background levels. Indel formation from Cas9 is estimated at ∼50% using ICE analysis. Inset, T7 endonuclease assay as a proxy for cutting efficiency demonstrates comparable frequencies of indel formation after transfection of either the I-SceI or Cas9 expression vector; however, note that this assay is much less sensitive than ICE for Cas9-generated indel formation, but ICE is not adaptable to I-SceI cleavage. n=2 experiments with 3 technical replicates for each timepoint. **B.** Quantification of colony formation. Sorted cells that have undergone DSB-induced LOH (dClover2+) form colonies with slightly reduced efficiency compared to sorted heterozygous cells or all cells in the population. No significant differences; Ordinary one-way ANOVA, with Tukey’s multiple comparisons test. n=2-4 independent sorting experiments, typically plating 1,000 and 2,000 cells for each, with plates from one experiment shown in Fig. 1E**,F**. **C.** Relative doubling time in proliferation assays. Doubling time, calculated by ln(2)/slope, is plotted normalized to the all cells condition. \**P*≤0.0129; Ordinary one-way ANOVA, with Tukey’s multiple comparisons test. n=3 experiments, 3-4 technical replicates per condition per experiment, with one experiment shown in Fig. 1G. **D.** Metaphase spread from Flo-LOH parental cells after FISH with a chr14 paint. While both chr14s are identified by the paint, the B6 chr14 can be differentiated from the 129 chr14 by the darker centromeric staining.

**Figure S3.**
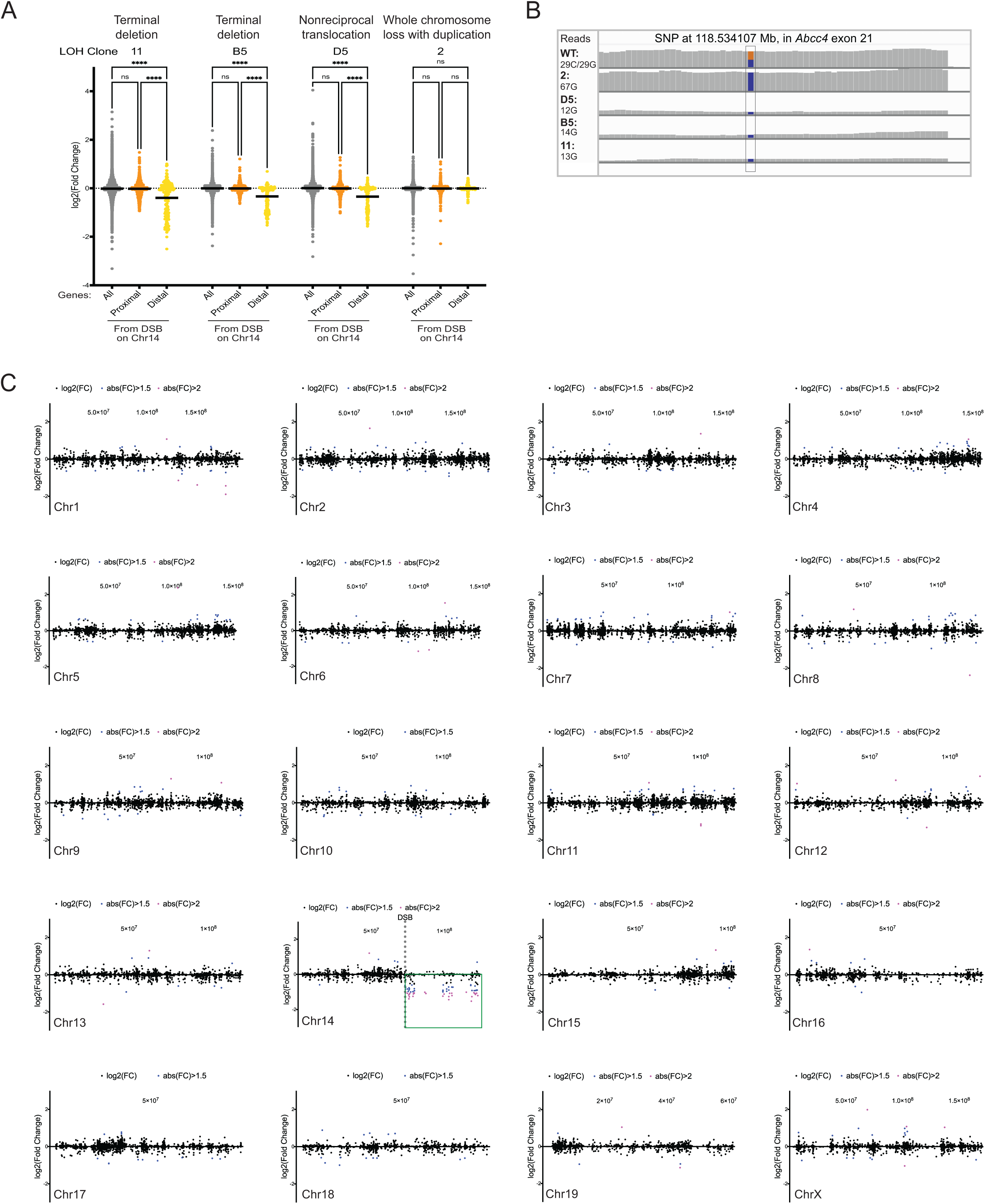
Only chromosome 14 which has undergone a DSB experienced a large regional decrease in gene expression. **A.** RNA-Seq analysis. The Log2 fold change compared to wild type is shown for all transcripts, transcripts from chr14 centromere-proximal to the DSB, and transcripts from chr14 distal to the DSB for each LOH clone. The three clones with copy number changes after the DSB exhibit a significant decrease in expression distal to the DSB, while the clone with copy neutral LOH is unchanged in expression of the region distal to the DSB. *\*\*\*\*P*<0.0001; Ordinary one-way ANOVA, with Tukey’s multiple comparisons test. **B.** Loss of transcripts occurs from the chr14 that experienced the DSB. Representative read counts from RNA-seq of a SNP in exon 21 of the *Abcc4* gene (rs237164364, location from GRCm38), located in the LOH region ∼45 Mb distal to the DSB on chr14. Half of the reads from the parental cells contain the SNP from the 129 chr14 (C) and half from the B6 chr14 (G). All four LOH clones only show the B6 SNP. The wild type and copy neutral clone had a similar number of reads, while the clones with segmental loss had fewer total reads. **C.** RNA-seq analysis for each chromosome from terminal deletion clone B5 relative to wild-type Flo-LOH cells. Only chr14 genes distal to the DSB (green box) show a consistent decrease in gene expression.

**Figure S4.**
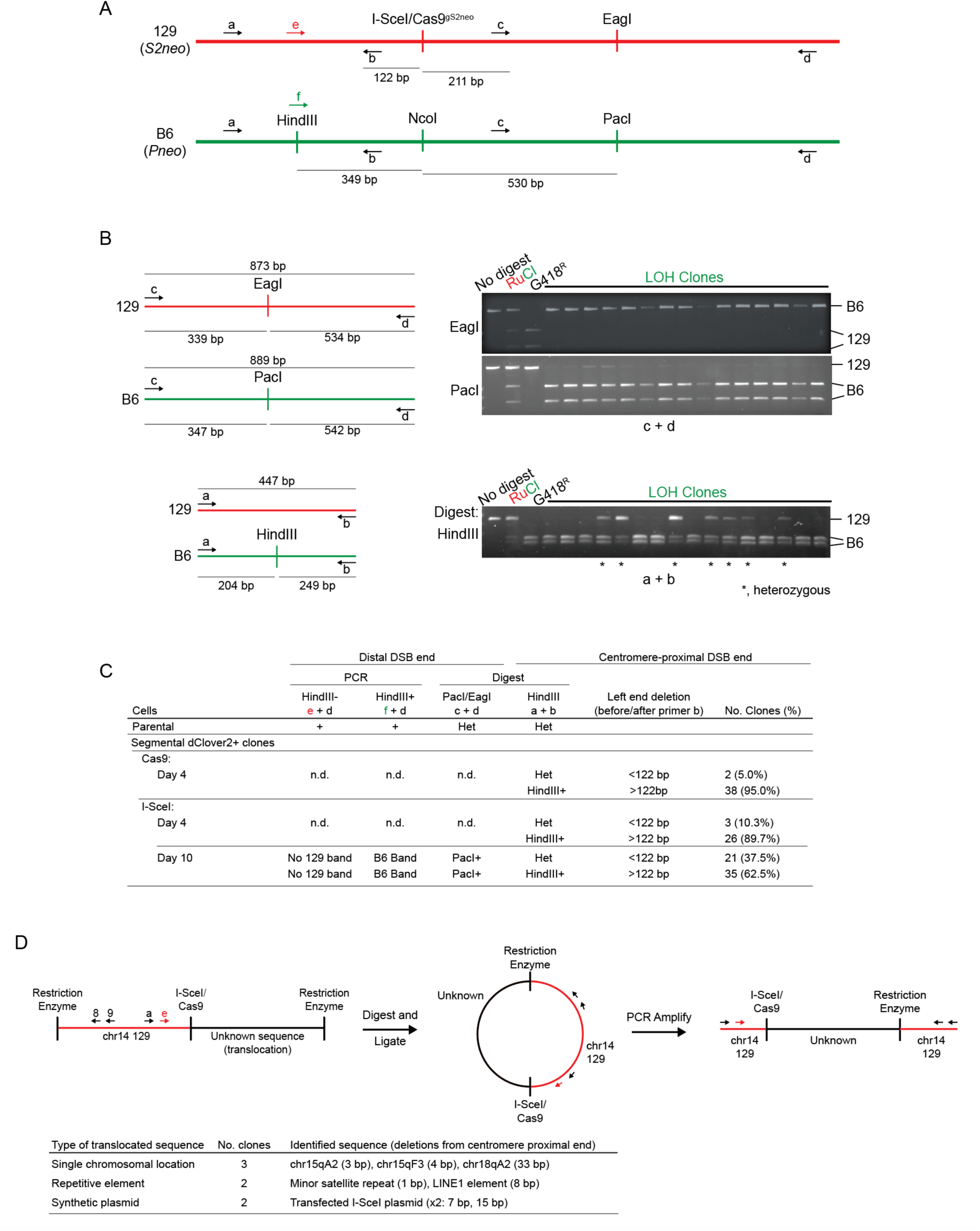
Characterization of clones with segmental loss. **A.** Polymorphisms between the 129 and B6 chromosomes near the DSB site in Flo-LOH cells. DSBs are introduced by either I-SceI or Cas9 which have overlapping recognition sites within a mutant *neo (S2neo)* gene integrated on the 129 chr14. The B6 chr14 has an NcoI site at this position within the *Pneo* gene^36^. Relative locations of universal primers that recognize either allele are represented by black arrows and allele-specific primers by red (e, 129 allele) and green (f, B6 allele) arrows. For orientation, the chromosomal coordinate of the first base of primer a is 14:73,450,528 (GRCm39, B6). **B.** Representative images of restriction digests of the indicated amplicons. All dClover2+ LOH clones have maintained B6 alleles, as evidenced by HindIII (a and b primers) and PacI (c and d primers) cleavage and have lost the 129 chr14 distal amplicon (c and d primers), since EagI cleavage is not observed. However, some of the clones have maintained heterozygosity at the centromere-proximal HindIII site, as evidence by the presence of an uncleaved amplicon from the a and b primers. LOH clones analyzed in the gels shown here are derived from the day 10 sort of dClover2+ cells after transfection of the I-SceI expression vector. RuCl, DNA from Flo-LOH parental cells; G418^R^, a G418-selected *neo+* IH-HR control clone which has the EagI site but not the PacI site. **C.** The centromere-proximal end of the DSB is better preserved in the single-positive dClover2 clones derived from the day 10 sort compared with those derived from the day 4 sorts. The first three columns in the table test for LOH distal from the DSB site. Allele-specific e or f forward primers were used at the HindIII polymorphism together with the reverse universal d primer distal to the DSB, as shown in **A**. While the B6 amplicon was found in all clones, the 129 amplicon was absent from all clones due to loss of the primer d binding site. Similarly, the universal c and d primers amplify distal to the DSB site on the B6 chr14, giving rise to a PacI cleavable amplicon, whereas this EagI cleavable amplicon is absent due to LOH in all clones. The fourth column tests for integrity of the centromere-proximal DSB end. All amplicons from primers a and b maintain the B6 chr14 HindIII+ polymorphism, while only some contain the 129 chr14 HindIII- polymorphism. Amplification around the HindIII sites requires the presence of the b primer binding sequence, located 122 bp upstream of the DSB site. Those without the amplicon from the 129 chr14 have segmental loss that extends >122 bp. In some clones LOH likely extends megabases, as with clones derived from the *Pneo* gRNA (Fig. 3E). Since endonuclease expression is transient, cells undergo LOH early in the experiment, but they can be out competed over time, suggesting that those with greater loss from the DNA end by day 10 have a significant growth disadvantage. **D.** Identification of translocated sequences in LOH clones using inverse PCR. Four restriction enzymes with recognition sequences proximal to the DSB (AflII, ScaI, PsiI, and NsiI) were used separately to digest genomic DNA from dClover2+ clones, which was then circularized by ligation. The translocated sequences were amplified using nested primers from the adjacent 129 chr14 sequences, and then identified by sequencing. Use of primer a and primer e restricted this approach to those clones that maintained the 129 chr14 HindIII polymorphism. Translocated sequences were identified in 7 clones and were derived from various sources. Deletions from the centromere-proximal end of the DSB were small, ranging from 1-33 bp.

**Figure S5.**
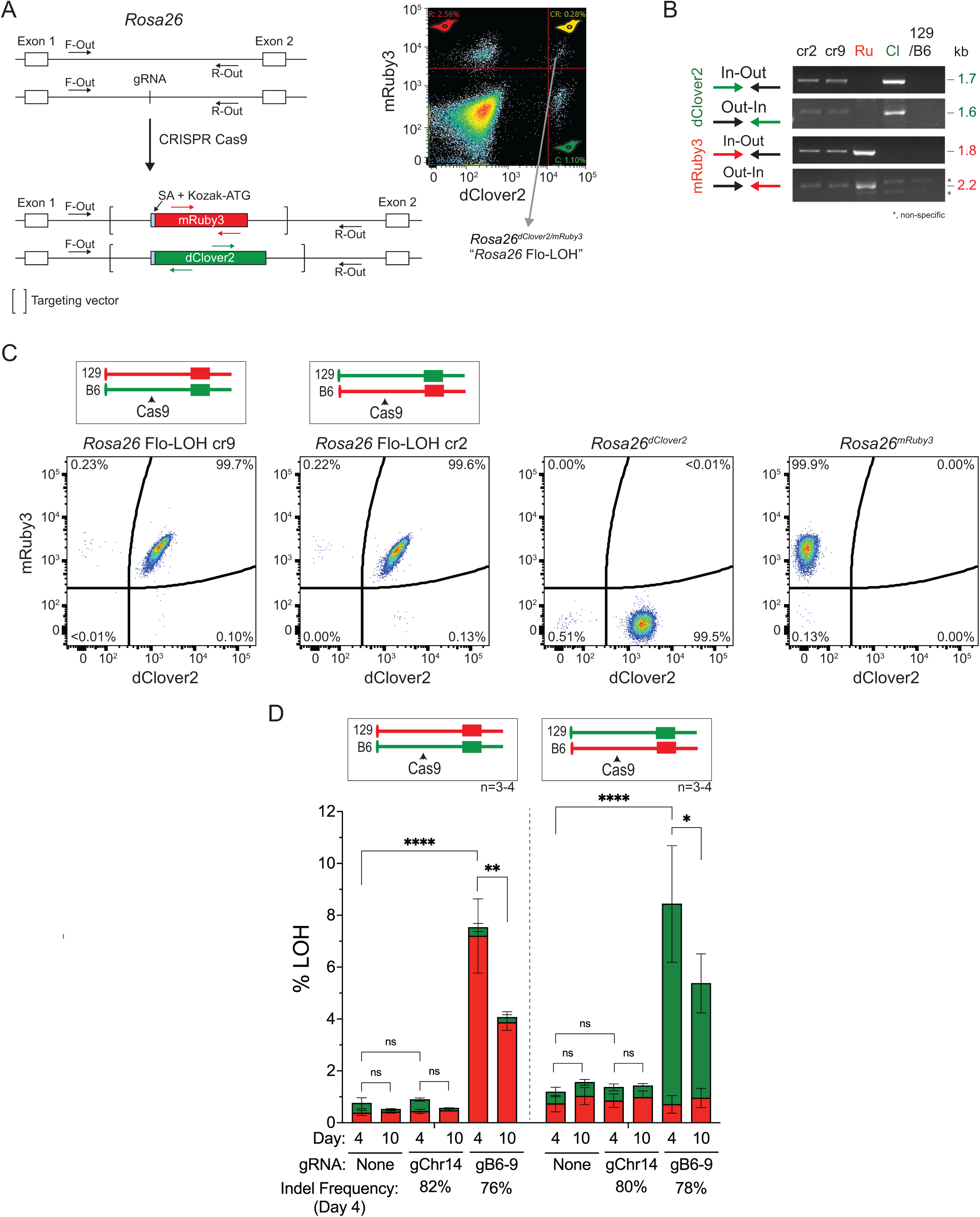
Flo-LOH: Targeting dClover2 and mRuby3 to the *Rosa26* locus in 129/B6 mESCs. **A.** mESCs were transfected with a Cas9 and gRNA expression vector and *Rosa26* targeting vectors with *mRuby3* and *dClover2* coding sequences. The fluorescent markers contain a splice acceptor and a Kozak sequence and become expressed upon correct integration in *Rosa26* intron 1. The brackets delineate the extent of the *Rosa26* homology arms in the targeting vectors. Double-positive cells were sorted from the transfected population to single cells by flow cytometry to generate *Rosa26* Flo-LOH cells (*Rosa26^dClover2/mRuby3^*). Single-positive cells were also sorted to generate compensation controls for subsequent flow cytometry. **B.** PCR confirmation of correct dual integration at the *Rosa26* locus. Primer locations are shown in **A**. Out-in and in-out primers use one primer outside the targeting vector and another primer within the fluorescent marker, as indicated. Both orientations were recovered: clone cr2 has integration of *dClover2* on the chr14 derived from the 129 mouse strain and *mRuby3* on the chr14 derived from the B6 strain (*dClover2*^129^;*mRuby3*^B6^), while clone cr9 has the opposite orientation (*mRuby3*^129^*;dClover2*^B6^). 129/B6, parental F1 hybrid cells. **C.** *Rosa26* Flo-LOH clones in both orientations uniformly express dClover2 and mRuby3, as seen by flow cytometry, with minimal spontaneous LOH (0.3-0.4%). Cells expressing only dClover2 or mRuby3 were used for controls in PCR and flow cytometry experiments. **D.** LOH from DSBs are chromosome specific. LOH on chr6 is induced by a DSB on chr6, but not on chr14. The percentage of cells that retain only dClover2 and mRuby3 expression is indicated by the green and red filled bars, respectively. Indel frequency was determined from the average of ICE analyses on the total cell population from all experiments. Error bars, mean ± s.d.; \*\*\*\**P*<0.0001, *\*\*P*=0.0038, *\*P*=0.0151 for total LOH; Ordinary one-way ANOVA, with Tukey’s multiple comparisons test.

**Figure S6.**
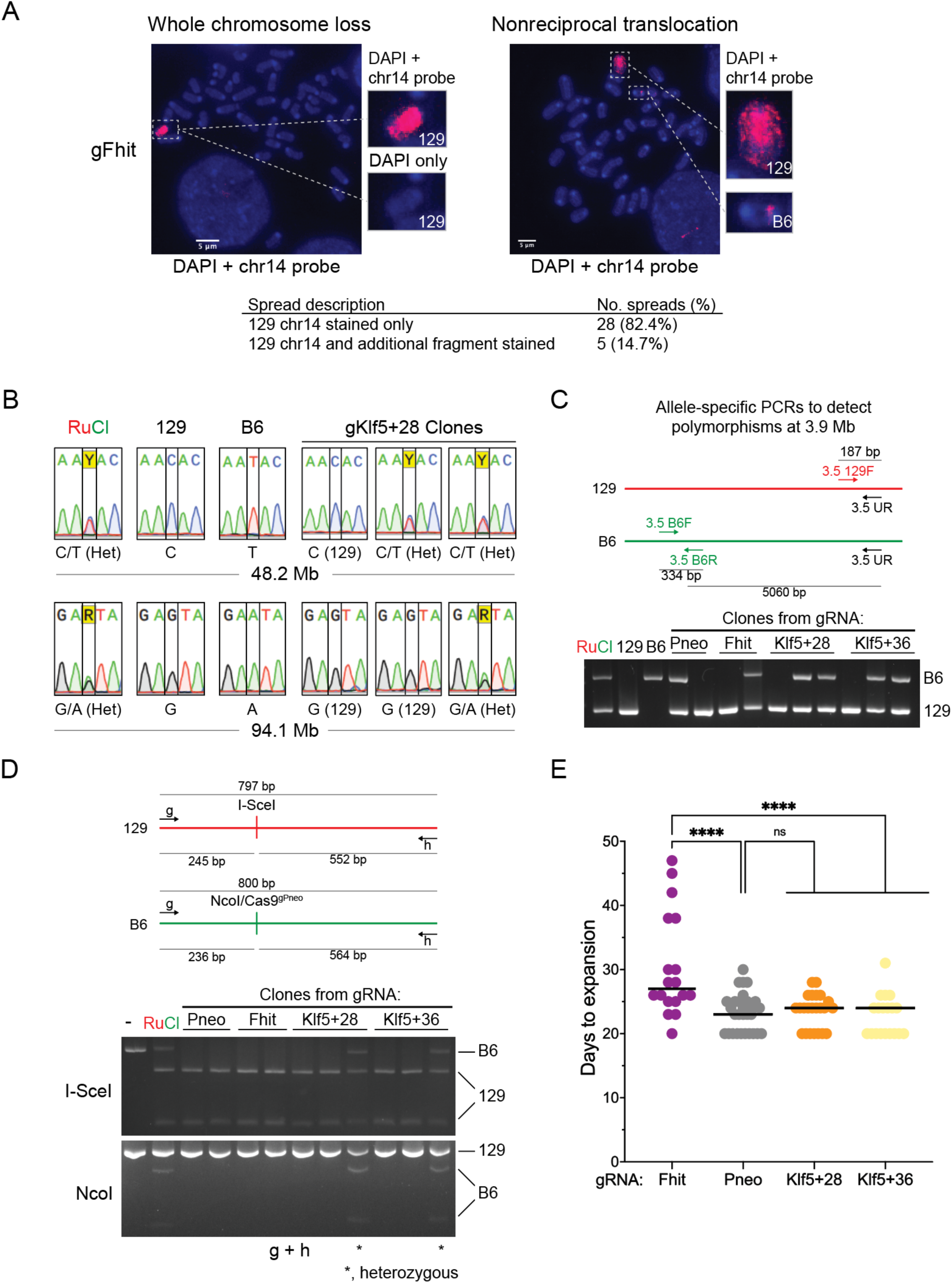
LOH can extend megabases centromere-proximal from the DSB site. **A.** Chromosome aberrations following a Cas9 DSB with gFhit at position 11.3 Mb on B6 chr14. (See also Fig. 3D.) After DSB induction by Cas9, single-positive mRuby3 cell populations were sorted, and metaphase spreads were stained with DAPI and probed with a whole chr14 probe. Top, representative images. In both images, an intact 129 chr14 is identified using a chr14 probe and lack of strong centromeric DAPI staining on the labeled chr14. In the left image, no additional chr14 staining is observed, indicating complete loss of the B6 chr14. In the right image, a nonreciprocal translocation is observed with a small region of the B6 chr14 fused with another chromosome or chromosomal fragment. Bottom, summary of spreads. The majority (>80%) of LOH cells lost the entire B6 chr14 or retained only an undetectable fragment of it, while ∼15% had some staining of an additional chromosome fragment, indicating a terminal deletion or nonreciprocal translocation involving the B6 chr14. **B.** SNPs at 48.2 Mb and 94.1 Mb in representative mRuby3+ LOH clones after a Cas9 DSB with gKlf5+28 at position 99.6 Mb on the B6 chr14. SNPs were amplified with universal primers and then sequenced to determine if heterozygosity was maintained or if LOH occurred. LOH in each case involved loss of the SNP from the B6 chr14, which had undergone the DSB. RuCl, DNA from Flo-LOH parental cells; 129 and B6, DNA from 129 cells and B6 cells, respectively. **C.** Polymorphisms at 3.9 Mb using allele-specific primers in representative mRuby3+ LOH clones generated after a DSB on B6 chr14 by the indicated gRNAs. For each gRNA, one clone is shown which only has the 129 polymorphism, consistent with loss of the broken B6 chr14, while the other clones maintain both the 129 and B6 polymorphisms, consistent with segmental loss (or interstitial loss). RuCl, DNA from Flo-LOH parental cells; 129 and B6, DNA from 129 cells and B6 cells, respectively. **D.** Polymorphisms in the mutant *neo* genes located at 73.5 Mb in representative mRuby3+ LOH clones generated after a DSB on B6 chr14. Universal primers amplify the NcoI site on the B6 chr14, which is adjacent to the Cas9 cleavage site, and the uncut I-SceI site on 129 chr14. Clones that are heterozygous have amplicons cleaved by both enzymes, while those that have undergone LOH have amplicons that are only cleaved by I-SceI. **E.** Days to expansion for mRuby3+ LOH clones after DSB formation from Cas9 and the indicated gRNAs, as a proxy for relative cell proliferation rates. It refers to the time required for a single cell to expand to sufficiently extract genomic DNA, freeze, and run flow cytometry. Clones targeted at *Fhit* (11.3 Mb) required significantly more time to fully expand than those targeted at *Pneo* (73.5 Mb) locus or 28 or 36 kb downstream of *Klf5* (99.6 Mb). \*\*\*\**P*<0.0001; Ordinary one-way ANOVA, with Tukey’s multiple comparisons test.

**Figure S7.**
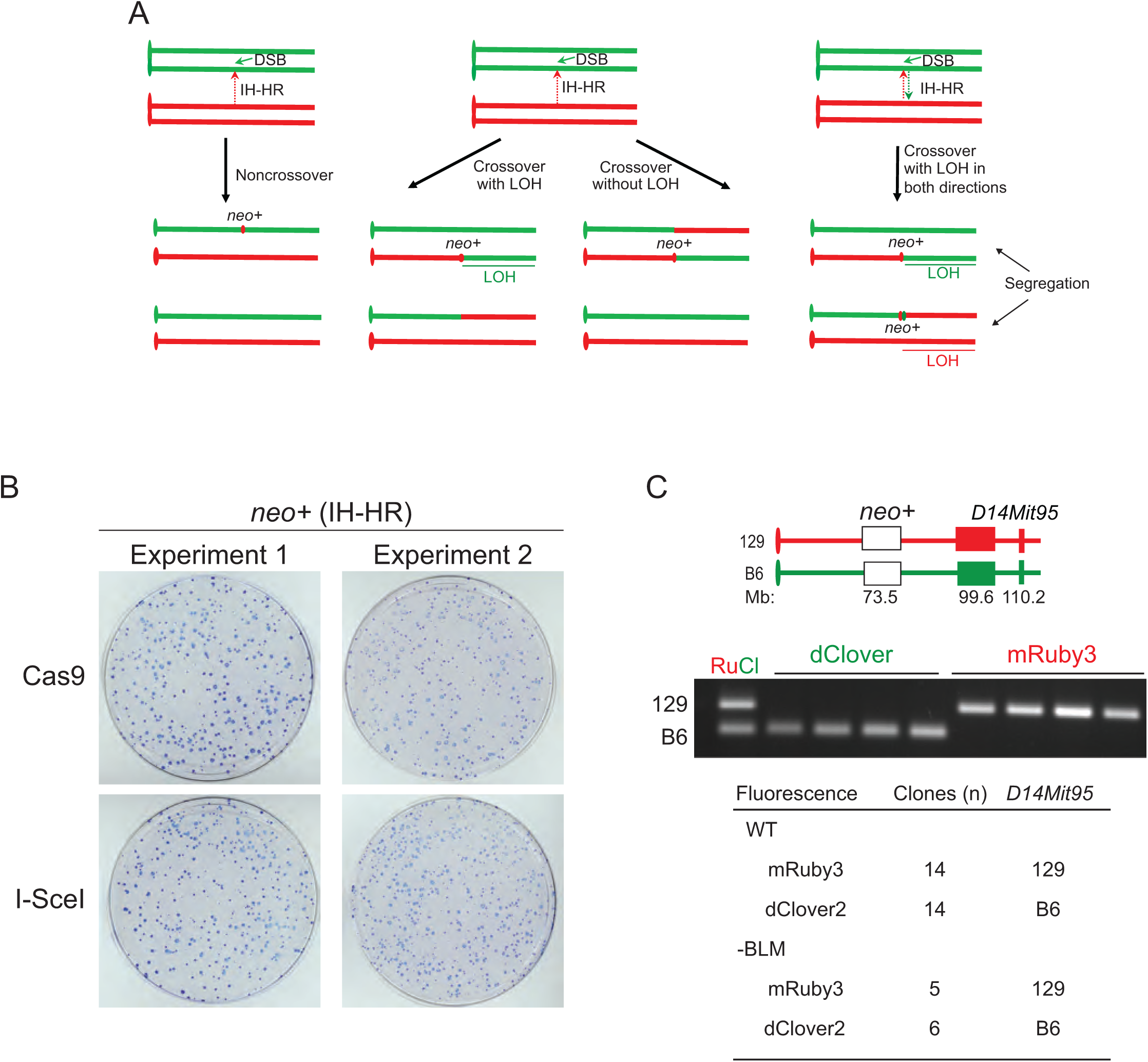
LOH in *neo+* cells after IH-HR. **A.** IH-HR outcomes leading to *neo+* cells. In noncrossovers, DSB repair incorporates homologous sequences from the chromosome homolog at the site of the DSB, leading to a small segment of LOH at the *neo+* gene. However, in crossovers, LOH extends from the DSB site to the distal telomere if the exchange chromosomes segregate away from each other in daughter cells. More complex events are also possible, as for example in the third scenario in which sequence information is transferred in both directions during IH-HR to create *neo+* genes on both chromatids; with the indicated segregation, LOH occurs in both daughter cells in opposite directions. **B.** IH-HR frequency is similar after induction of a Cas9 or I-SceI DSB at *S2neo*. Flo-LOH cells were transfected with a Cas9 or I-SceI expression vector and plated. To select for *neo+* colonies, G418 was added 24 hr later, and replaced every 3-4 days until colonies formed. One representative plate for Cas9 and I-SceI is shown from two separate experiments in which 0.9 million cells were plated. **C.** Distal LOH from DSB-induced IH-HR is confirmed in wild-type and BLM-depleted cells by PCR analysis of the *D14Mit95* locus in *neo+* LOH clones.

**Figure S8.**
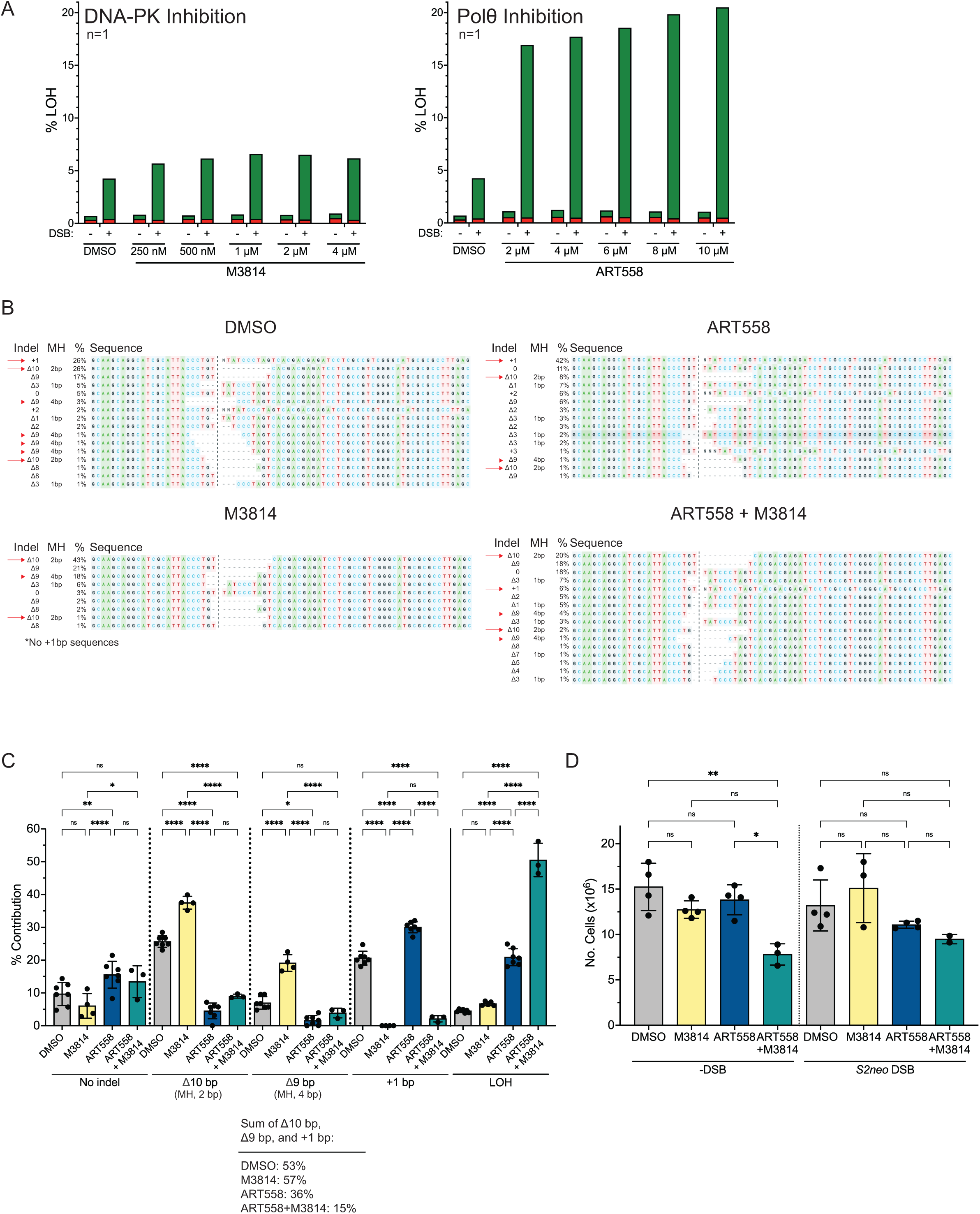
Inhibition of Pol 8 and DNA-PK impacts DSB repair events at the *S2neo* site. **A.** Titration of end joining inhibitors M3814 (DNA-PK) and ART558 (Pol θ) for their effects on LOH. At varying concentrations, both inhibitors increase in LOH after DSB induction (*S2neo* gRNA), although the effect with M3814 is marginal, especially compared with Pol 8 inhibition, which drastically increases LOH. Given the small dose-dependent increase in DSB-induced LOH with ART558, 6 μM was chosen for further experiments. Since LOH plateaus with M3814 at 1 μM, this concentration was chosen for further experiments. **B.** Sequences of indels after DSB repair at the *S2neo* DSB in the presence of end joining inhibitors from ICE analysis. The two main end joining products (+1 and Δ10, red arrows) are conversely affected by M3814 and ART588, consistent with inhibition of NHEJ and MMEJ, respectively, such that the +1 product is abrogated by M3814 but increased with ART558, while the Δ10 product is increased with M3814 but decreases with ART588. Note the presence of the 2 bp GT microhomology (MH) in the Δ10 product and that imperfect MH extends further upstream (CCCT). A Δ9 bp product at a 4 bp perfect homology (CCCT, red arrowhead) also increases with M3814 and is suppressed by ART558, consistent with that product forming preferentially by MMEJ. Note that with the Δ10 and Δ9 products, the ICE sequencing results place the microhomology in different locations for the same product, hence the multiple arrows and arrowheads, respectively. Results are shown from one experiment; a summary of results from all experiments (n≥3) is in **Fig. S8C**. **C.** Comparison of DSB outcomes at the *S2neo* site with Pol 8 and/or DNA-PK inhibition. LOH frequencies are from **Fig 5E**. The frequencies of no indel, Δ10 bp with 2 bp microhomology, Δ9 with 4 bp microhomology, and +1 events are from ICE sequence contribution (**Fig. S8B** and replicates) and are normalized to adjust for LOH frequencies. For example, with 20% LOH, the end-joining percentage is reduced by 20%. Note that the unmodified sequences (0 bp class) are reduced with M3814, possibly due to inhibited precise NHEJ, while this 0 bp class tends to be increased with ART558. \*\*\*\**P*<0.0001, \*\**P*=0.0041, \**P*≤0.0239; Ordinary one-way ANOVA, with Tukey’s multiple comparisons test. **D.** Effect of inhibitors on cell proliferation. Pol 8 inhibition (ART558) or DNA-PK inhibition (M3814) has little effect on cell proliferation either with or without a DSB at *S2neo*, while combined inhibition leads to a more discernable effect. After transfection of the *S2neo* gRNA, 200,000 cells were plated in a 10-cm plate and then counted 4 days later. \*\**P*=0.0031, \**P*=0.0217; Ordinary one-way ANOVA, with Tukey’s multiple comparisons test. n=2-4

**Figure S9.**
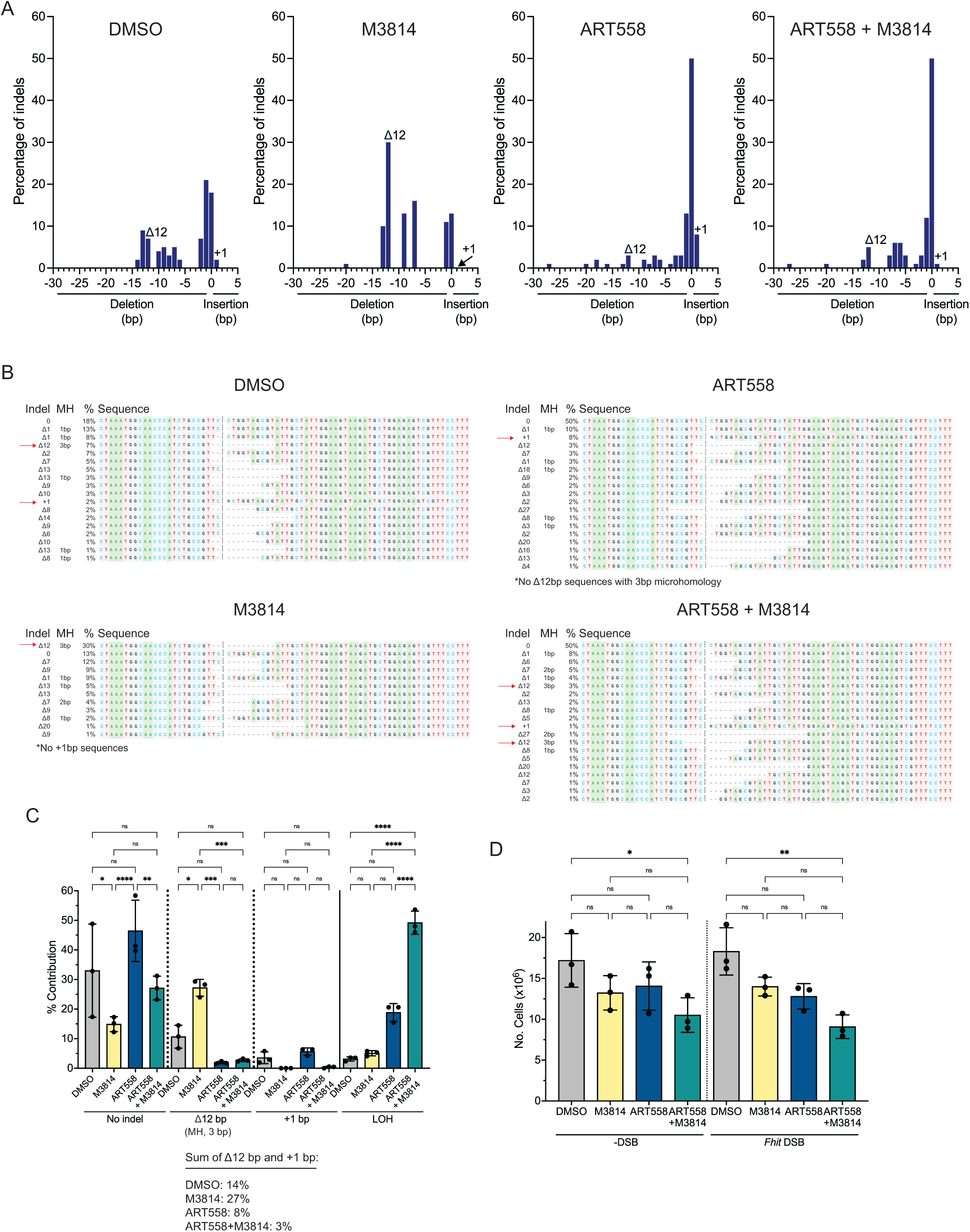
Inhibition of Pol 8 and DNA-PK impacts DSB repair events at the *Fhit* site. **A.** Indel distribution after DSB repair in the presence of end joining inhibitors using ICE analysis. Two end joining products at the *Fhit* DSB, +1 and Δ12 with 3 bp microhomology, are conversely affected by M3814 and ART588, consistent with inhibition of NHEJ and MMEJ, respectively. Results are shown from one experiment; a summary of results for these indels from all experiments (n=3) is in **C**. **B.** Sequences of indels after DSB repair at the *Fhit* DSB in the presence of end joining inhibitors from ICE analysis. The two main end joining products (+1 and Δ12, red arrows) are differentially impacted by M3814 and ART588. The +1 product is abrogated by M3814 but increased with ART558, while the Δ12 product is increased with M3814 but abrogated with ART588. The sequences are from the same experiment shown in **A**. **C.** Comparison of DSB outcomes at the *Fhit* site with Pol 8 and/or DNA-PK inhibition. LOH frequencies are from **Fig 5E**. The frequencies of no indel, Δ12 bp with 3 bp microhomology, and +1 events are from ICE sequence contribution (**B** and replicates) and are normalized to account for LOH frequencies, which comprise a large part of the total DSB outcomes. \*\*\*\**P*<0.0001, \*\*\**P*≤0.0002, \*\**P*=0.0059, \**P*<0.0383; Ordinary one-way ANOVA, with Tukey’s multiple comparisons test. **D.** Effect of inhibitors on cell proliferation. Pol 8 inhibition (ART558) or DNA-PK inhibition (M3814) has a small but insignificant effect on cell proliferation either with or without a DSB at *Fhit*, while combined inhibition leads to a significant reduction in cell number. After transfection of the *Fhit* gRNA, 200,000 cells were plated in a 10- cm plate and then counted 4 days later. \*\**P*=0.0031, \**P*=0.0421; Ordinary one-way ANOVA, with Tukey’s multiple comparisons test. n=3

